# Thrombin-free polymerization leads to pure fibrin(ogen) materials with extended processing capacity

**DOI:** 10.1101/2020.05.12.091793

**Authors:** Clément Rieu, Gervaise Mosser, Bernard Haye, Nicolas Sanson, Thibaud Coradin, Léa Trichet

## Abstract

Fibrin is a key protein for various clinical applications such as tissue reconstruction. However, in contrast to type I collagen, fibrin shaping has so far faced major limitations related to the necessity to add thrombin enzyme to fibrinogen precursors to induce fibrin self-assembly. Here we report a thrombin-free gelation pathway of fibrinogen solutions by incubation at 37°C in mild acidic conditions. We unravel the biochemical mechanisms underlying the gelation process and draw comparison between fibrinogen and fibrin at both molecular and supramolecular levels in these conditions. The protocol enables to control the viscosity of fibrin(ogen) solutions, and to induce fibrin(ogen) gel formation by simple 37°C incubation, with a reinforcement effect at neutralization. It facilitates processing of fibrin(ogen) materials, for coating, molding and extrusion, and offers new possibilities such as 3D printing. This approach is further compatible with type I collagen processing and can provide advanced tissue engineering scaffolds with high bioactivity.

## Introduction

Fibrin is a gold standard material in tissue engineering ^1,2^. It exhibits high bioactivity thanks to the direct adhesion sites for integrins ^3,4^, which impact cell migration, proliferation and fate ^5,6^. Fibrin also indirectly interacts with cells through binding to extracellular matrix proteins ^7^, polysaccharides ^8^ and growth factors ^9^. It is widely used clinically, mainly as a hemostat, sealant and adhesive. *In vivo*, fibrin is a major actor of haemostasis ^10^, which is induced by a complex reaction cascade initiated by the adhesion of platelets at the site of injury ^11,12^. A key step is the thrombin mediated conversion of circulating fibrinogen into fibrin, which subsequently polymerizes to form a tight plug ^13^.

Fibrinogen is a 340 kDa protein made of three pairs of chains, called α, β and γ that assemble in a specific symmetric shape, forming 3 aligned globules linked together by coiled coil connectors constituted by the three different chains ^14^ (Fig.1.A.1). The central domain, called E, gathers the amino-terminal parts of the 6 chains, linked together by di-sulfide bonds. The distal domains, called D, are made of the carboxy-terminal ends of the β and γ chains. The carboxy-terminal ends of α chains, called αCs, further extend in a random-coiled manner, with globular domains at their ends. Those domains interact together and with the E domain, and hence do not float freely. During clotting, thrombin cleaves two pairs of fibrinopeptides FpA and FpB from the central E nodule, (Fig.1.A.2 **(0)**, where only two out of four Fps are drawn for clarity purpose) ^15^. This cleavage reveals two sites A and B which bind to sites a and b, respectively located on the γ and β chains in the D nodule of an adjacent molecule. Such interactions promote the polymerization of fibrin monomers in a half staggered manner (Fig.1.A.2 **(0’)**), resulting in gels (Fb_N_) (Fig.1.A.3) with a highly fibrillar network (Fig.1.A.4). A certain delay, called clotting time, exists before the formation of a macroscopic clot ^13^. Lateral assembly of protofibrils only occurs at the end of the clotting time, which can last from seconds to hours depending on pH and ionic strength ^16^.

**Fig. 1.**
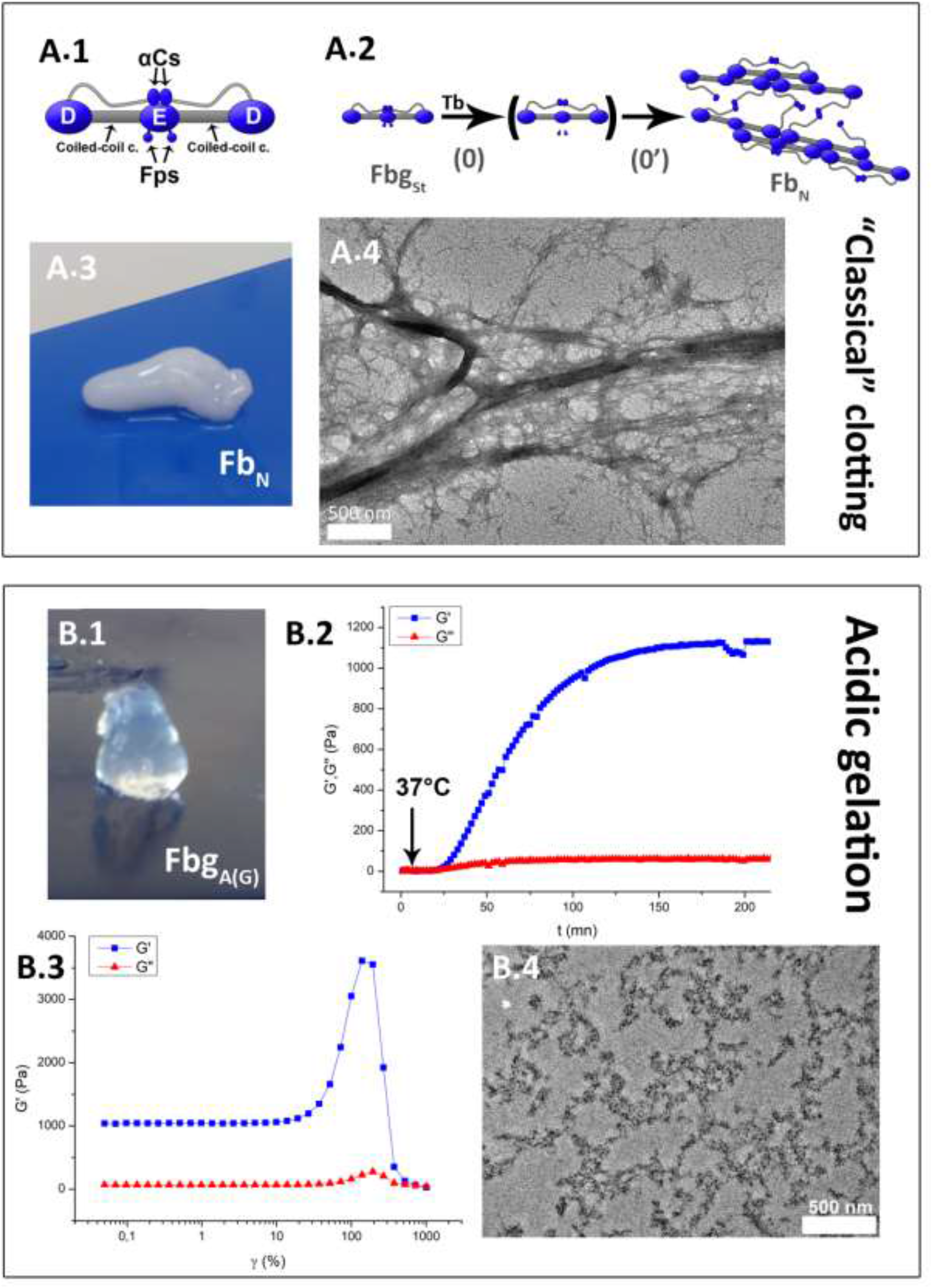
Two gel formation mechanisms: “classical” neutral thrombin-driven (A) and acidic thrombin-free (B). (A.1) Schematic structure of fibrinogen, made of two distal D domains and one central E domain, linked by coiled-coil connectors. The E domain bears the two pairs of fibrinopeptides (Fps) A and B. The two carboxy-terminal ends of the α-chains further extend and fold into small globular domains, noted αCs, which connect to the central E domain. (A.2) “Classical” mechanism where the fibrinopeptides are cleaved **(0)** by thrombin (Tb), converting the fibrinogen proteins (Fbg_St_) into fibrin monomers that self-assemble **(0’)** in a half-staggered manner to form a fibrin clot (Fb_N_). (A.3) Image of a fibrin clot. (A.4) Fb_N_ observed by Transmission Electron Microscopy (TEM). (B.1) Image of an acidic gel (Fbg_A(G)_) formed by incubation of a 40 mg/mL fibrinogen solution at pH 3.6 (Fbg_A_) at 37°C. (B.2) Rheological dynamical analysis of gel formation of Fbg_A(G)_ initiated by a temperature switch from 20°C to 37°C. (B.3) Mechanical behavior of Fbg_A(G)_ under amplitude sweep. (B.4) Structure of an acidic gel Fbg_A(G)_ included in araldite and observed by TEM.

Difficulty to shape natural materials into 3D structures remains a major obstacle to mimic cell environment ^17^. The necessity to mix fibrinogen and thrombin to trigger self-assembly and the involved delay may account for the scarcity of work on fibrin shaping into 3D structures. Fibrin is mostly used in the form of gels that can be molded ^18^, or in addition to other materials ^19^. Some attempts have been carried out to produce fibrin microbeads ^20^ or nanostructures ^21,22^. So far, only one process has been reported for pure fibrin extrusion ^23^, where fibrinogen and thrombin kept in different solutions are mixed together with a Y connector before extrusion. In sharp contrast, collagen, another gold standard material in tissue engineering, has been extensively studied ^24^. The simplicity of collagen self-assembly, induced by neutralization or 37°C heat activation, makes it favorable for multiple shaping techniques, such as molding or extrusion ^25^. To investigate if similar processes could be used to trigger fibrin polymerization and thus widen the possibilities in fibrin shaping, we thus focused on fibrinogen and fibrin solutions at acidic pH.

### Acidic fibrinogen gel formation

Stock solutions of fibrinogen (Fbg_St_) at 40 mg/mL were acidified to pH 3.6, the highest pH below the isoelectric point (~5.5) at which the solution remained stable at ambient temperature (20°C) without precipitating. Without any addition of thrombin these acidic solutions (Fbg_A_) incubated at 37°C formed within an hour a transparent brittle gel (Fbg_A(G)_) (Fig. 1.B.1), while solutions left at ambient temperature set to a gel only after several weeks. Gel formation kinetics was increased at lower pH, with instantaneous gelation at pH 2.5. Formation of fibrinogen transparent gels at acidic pH had already been briefly evoked by Fay & Hendrix ^26^ and Mihalyi ^27^. However no study of this phenomenon has been carried out so far.

Gel formation from acidic solutions at pH 3.6 was dynamically investigated by switching the temperature of geometries from 20°C to 37°C after 5 minutes of rheological properties measurement (Fig. 1.B.2). For approximately 15 minutes G’ remained constant, and then rose to reach approximately 1 kPa after three hours, overtaking G” (factor > 10). During frequency sweep on the newly formed gels Fbg_A(G)_, G’ increased with frequency, with an inflection point at 1.6 rad/s, while G” exhibited a reverse bell shape (SI Appendix, Fig. S1). During amplitude sweep, gels exhibited first a plateau with G’ around 1 kPa till around 20 % amplitude, then a 3 to 4-fold strain stiffening until 130% strain (Fig. 1.B.3). As a comparison Wufsus *et al.* reported a G’ value one order of magnitude higher for “classical” Fb_N_ clots made from 30 mg/mL and 50 mg/mL initial fibrinogen solutions ^28^. Structure-mechanics relationship in “classical” fibrin clots have been widely investigated but no general picture has been proposed yet. Non-linear elasticity of fibrin clots Fb_N_ originates from the multi-scale hierarchical structure ^29,30^. The observed behavior in frequency sweep of Fbg_A(G)_ differs from that of classical clots and exhibits a signature reminiscent of rubber plateau in polymer melts with high-molecular-weight entangled chains ^31^. However, the strain-stiffening behavior is characteristic of biological gels and in particular of fibrin ^29^. AFM force-unfolding experiments on fibrin proteins suggest that clot elasticity may arise from coiled-coil and γ-nodule unfolding, and from αCs interactions ^32,33^. Coiled-coil and/or γ nodule unfolding might then account for elasticity and strain stiffening behavior of the acidic gels, while it is unlikely that αCs are involved as they do not interact at acidic pH ^34^.

Transmission Electron Microscopy (TEM) observations of sections of fibrinogen gels Fbg_A(G)_ included in araldite display a network of fine entities without any specific pattern (Fig. 1.B.4). After chopping a gel in water and depositing the supernatant on a carbon coated grid, some large fiber-like structures can be observed (Fig. 1.B.5). In comparison, a “classical” fibrin clot made by mixing fibrinogen and thrombin at neutral pH exhibits a highly fibrillar network with fibers of various sizes (Fig. 1.A.4).

### Underlying Mechanism of acidic gel formation

Acidic fibrinogen gels Fbg_A(G)_ were surprisingly stable to most conditions known to disperse protein-based hydrogels (SI Appendix, Table S1). Change to neutral or basic pH did not dissolve the gels. They were resistant to low and high ionic strength, as well as to high temperature and salt-in solutions. Gels were however soluble in urea. As expected they could also be dissolved in DTT, since the constituting fibrinogen chains are linked together by multiple disulfide bonds. SDS electrophoretic gels of acidic fibrinogen gels dissolved in urea confirmed that no major cleavage was provoked by the acidic conditions and by the thermal treatment at 37°C (SI Appendix, Fig. S2). MALDI-TOF performed on fibrinogen acidic gels Fbg_A(G)_ showed that no fibrinopeptide could be detected after acidic gel formation, while both Fps A and B were visible for fibrin clots FbN (SI Appendix, Fig. S3). Thus, the mechanism involved in gel formation at acidic pH is different from the “classical” one (**0+0’**, Fig. 1.A.2).

Circular dichroism measurements on fibrinogen stock Fbg_St_ and acidified solutions FbgA demonstrate a change in the secondary structure upon acidification, with a decrease in intensity of the 225 nm peak (Fig. 2.A). No further change in acidic solution signature could be observed upon incubation at 37°C, even after formation of a gel Fbg_A(G)_. The change appears to be reversible for the solutions Fbg_A_ when raising the pH above the pI (Fig. 2.A). CD spectra of Fbg_st_ have already been reported and a consensus of approximately a third of α-helix content and a third of random structure exists, while the content of the remaining structures fluctuates according to studies ^35–37^. In our case the change in structure upon acidification could not be reliably attributed. Similar spectra were however observed in different fibrinogen fragments ^37–39^ and partially heat-denatured fibrinogen ^35^, which tends to point out a modification of the helical structure and/or the partial denaturation of the D domain.

**Fig. 2.**
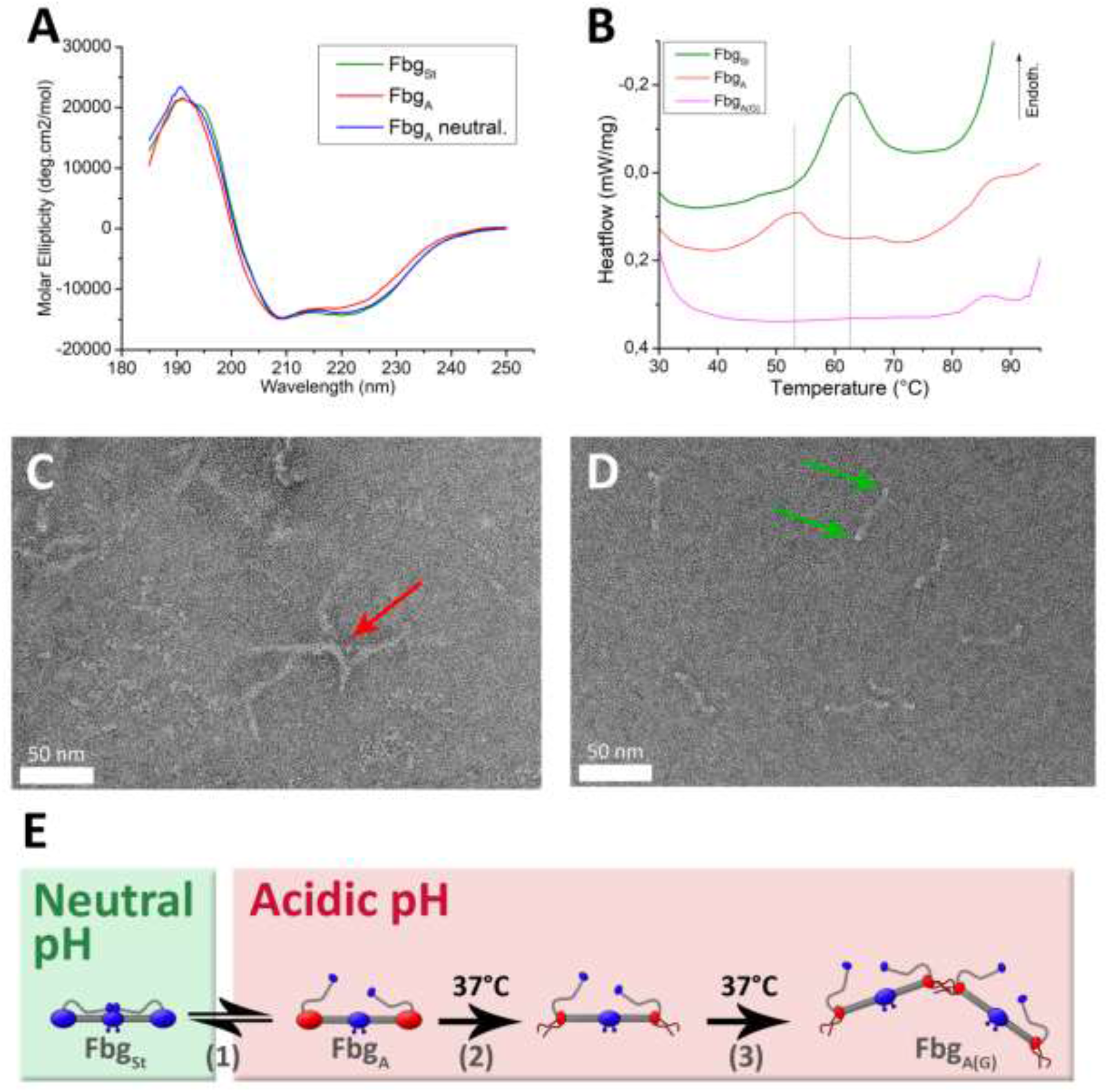
Mechanism of acidic fibrinogen gel formation. (A) Circular dichroism spectra at 20°C of a Fbg_st_ (green) solution, Fbg_A_ solution at pH 3.6 with HCl (red), and the same solution brought back to neutral pH by addition of NaOH (blue). (B) Thermograms of Fbg_st_ solution (green), Fbg_A_ solution pH 3.6 (red), and Fbg_A(G)_ gel pH 3.6. (C & D) Fbg_A_ and Fbg_st_, respectively, dried on a carbon coated grid and observed by TEM. (E) Schematic of the mechanism inferred from the obtained results. Upon acidification to pH 3.6 **(1)** of Fbg_St_, the D domains of acidic fibrinogen Fbg_A_ are destabilized in a reversible manner (red). Incubation at 37°C provokes the denaturation **(2)** of the thermolabile D domains of fibrinogen proteins that subsequently interact by their extremities **(3)** to give an acidic fibrinogen gel Fbg_A(G)_.

Differential Scanning Calorimetry of fibrinogen stock Fbg_St_ exhibits a major peak around 63°C (Fig. 2.B). This low temperature peak of fibrinogen at neutral pH is attributed to the D domain ^40^. When acidified, three peaks appear: a major peak around 52°C and a shoulder around 65°C, corresponding to the thermolabile part of the D domain and to the αCs, respectively ^40^, followed by a third peak close to 85°C, attributed to the E domain. The thermolabile D domain is thus destabilized by the acidic condition with a low temperature peak shift of −10°C, in accordance with previous results ^40^. When the acidic solution Fbg_A_ is first kept at 37°C for 1 hour before the scan, only the high temperature peak around 85°C remains (Fig. 2.B). The incubation of acidic solutions at 37°C for 1 hour thus leads to vanishing of the low temperature peak attributed to the thermolabile D domain ^40^. After incubation, the absence of the shoulder at 65°C, corresponding to the αCs and reported to be reversible upon cooling ^41^, could be due to a too low sensitivity of our setup. Observation of dilute acidic fibrinogen solutions Fbg_A_ by TEM shows that fibrinogen proteins tend to aggregate preferentially by their extremities (Fig. 2.C), on the opposite to fibrinogen proteins from stock solution Fbg_St_, which appear dispersed (Fig. 2.D).

Aggregation of fibrinogen in acidic conditions upon incubation in presence of D and E fragments was investigated by dynamic light scattering. Altered aggregation in presence of D fragments was observed, on the opposite to E fragments (Appendix, Fig. S4), which tends to confirm that D domains are involved in aggregation.

As summed up in the sketch (Fig. 2.E), stock fibrinogen Fbg_St_ undergoes upon acidification **(1)** a reversible change in structure that corresponds to a destabilization of its thermolabile D domain (represented by a change of color of the D domain from blue to red) in the acidic fibrinogen Fbg_A_. Upon incubation at 37°C **(2)**, these domains are denatured (represented by the floating carboxyterminal ends of β and γ chains ^42^), promoting interactions between proteins **(3)** and resulting in a gel Fbg_A(G)_.

### Effect of neutralization on fibrinogen and fibrin acidic gels

#### Fibrinogen

The acidic fibrinogen gels Fbg_A(G)_ are not soluble when dipped in a neutral solution and no major change in aspect such as opacification or gel contraction can be observed. The change in rheological properties upon neutralization of acidic gels Fbg_A(G)_ was studied by forming a gel at 37°C in acidic condition and then adding a neutral solution of 100 mM Hepes around the geometries (Fig. 3.A, blue). This resulted in a gel Fbg_N(G)_ with a two-fold increase in storage modulus as well as a different signature in frequency sweep which now increases linearly with the frequency (SI Appendix, Fig. S5). These modified mechanical properties may be related to a change in structure.

**Fig. 3.**
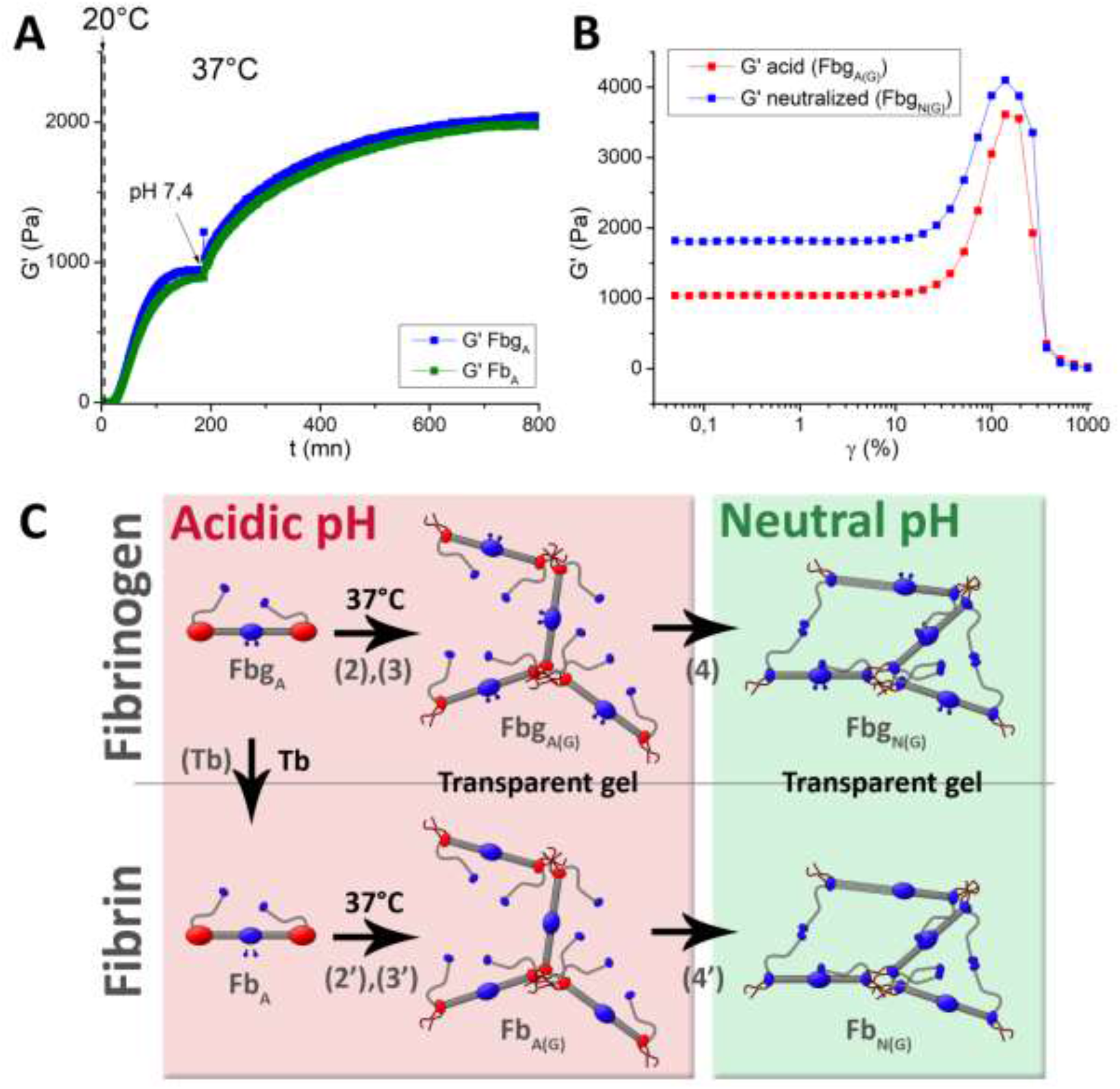
Effect of neutralization on acidic fibrin and fibrinogen gels. (A) Formation of acidic gels from acidic fibrinogen Fbg_A_ and fibrin Fb_A_ solutions by incubation at 37°C followed by gel stiffening upon addition of a neutral solution around the geometry. (B) Comparison of mechanical response in amplitude sweep of a neutralized acidic fibrinogen gel (red) compared to a non-neutralized one (blue). (C) Schematic of the underlying mechanisms inferred from the measures. Fibrinogen proteins Fbg_A_ assemble upon incubation **(2)** forming a gel Fbg_A(G)_ **(3)** as seen on Fig.2. Upon neutralization, creation of additional interactions inside the gel Fbg_N(G)_ lead to internal constraints and subsequent partial unfolding of some proteins **(4)**. Similar results are obtained in presence of thrombin (Fb_A_), both during the gel Fb_A(G)_ formation in acidic conditions at 37°C **(2’)(3’)** and upon neutralization **(4’)** to result in a similar gels Fb_N(G)_ as Fbg_N(G)_.

The increase in G’ and G” with frequency is closer to fibrin clot FbN signature ^28^. This modification is not surprising as protein global charge is modified secondary to pI crossing. In addition to this, the αCs may form intermolecular interactions. Veklich *et al.* reported that αCs interact together and also with the E domains at neutral pH, while they float freely in acidic conditions. They also demonstrated that αCs cleaved from fibrinogen were dispersed in acidic conditions and polymerized into fibers upon neutralization. It is then likely that the αCs float during gel formation at acidic pH and interact together upon neutralization. This would be facilitated by the forced proximity between proteins in the network. Regarding the amplitude sweep a similar behavior as in the acidic condition was observed (Fig. 3.B), with a G’ plateau for low amplitudes, at approximately twice the value, but only a slight increase of peak amplitude between 100 and 200 % strain. Such a phenomenon may be due to a partial initial stretching of the proteins inside the network upon neutralization (Fig. 3.C **(4)**) because of internal constraints related to electrostatic repulsion and/or αCs interactions in the neutral gels.

#### Fibrin

Till now, no thrombin was introduced in the solutions, gels being made of fibrinogen proteins (as confirmed by MALDI-TOF results). It was shown in literature that addition of thrombin to fibrinogen in acidic conditions effectively cleaves the Fps, transforming fibrinogen Fbg_A_ into fibrin monomer Fb_A_ but without inducing clotting ^43^, which becomes instantaneous upon neutralization. The influence of thrombin addition to our system was therefore investigated.

Fb_A(G)_ gels from acidic fibrin solutions Fb_A_ could be formed by incubation at 37°C, in the same fashion as Fbg_A(G)_, without any impact on kinetics or gel aspect. Same rheological study of acidic gel formation and subsequent neutralization was carried out from fibrin solution Fb_A_ as for fibrinogen Fbg_A_. No significant difference either in formation kinetics or mechanical properties could be observed between Fb_A(G)_ and Fbg_A(G)_, as well as between Fb_N(G)_ and Fbg_N(G)_ (Fig. 3.A). MALDI-TOF performed on a Fb_A(G)_ gel formed by incubation at 37°C confirmed that FpA and FpB are effectively cleaved upon addition of thrombin to acidic fibrinogen (SI Appendix, Fig. S3). Thus, the removal of the Fps by thrombin does not seem to affect the acidic gel formation and neutralization.

### Effect of neutralization on fibrinogen and fibrin viscous acidic solutions

#### Fibrinogen

As already mentioned viscosity increases progressively during incubation at 37°C (Fig. 1.B.2), thus giving the possibility to obtain viscous fibrinogen solutions Fbg_A(VS)_. When neutralized, these solutions form opaque gels Fbg_N(VS)_, stable over time, on the contrary to non-heated solutions Fbg_A_ which produce precipitates that dissolve over time. This phenomenon was investigated by examining rheological properties. Fbg_A(VS)_ heated prior to measurement were neutralized five minutes after starting the measurement at 25°C by careful addition of a neutral solution of 100 mM Hepes around the geometries, and resulted in gels Fbg_N(VS)_ with G’ of a few hundred of Pa after 24 hours (SI Appendix, Fig. S6). Solutions Fbg_A(VS)_ with different initial viscosities were analyzed and the G’ of the final gels Fbg_N(VS)_ were plotted as a function of the initial G’ of the solutions (Fig. 4.A.1). A sharp increase of final G’ as a function of initial G’ is observed, followed by a plateau around 350 Pa. Resulting gels Fbg_N(VS)_ exhibit dense regions of aggregated proteins, reminiscent of the structure observed in acidic fibrinogen gels Fbg_A(G)_ as observed by TEM (Fig. 4.A.2). It is likely that the mechanism **(5)** responsible for formation of the gels Fbg_N(VS)_ upon neutralization of Fbg_A(VS)_ is similar to the mechanism causing acidic gel Fbg_A(G)_ strengthening after raising pH (Fig. 3.C **(4)**), and was thus hypothesized to involve αCs interaction (Fig. 4.A.3).

**Fig. 4.**
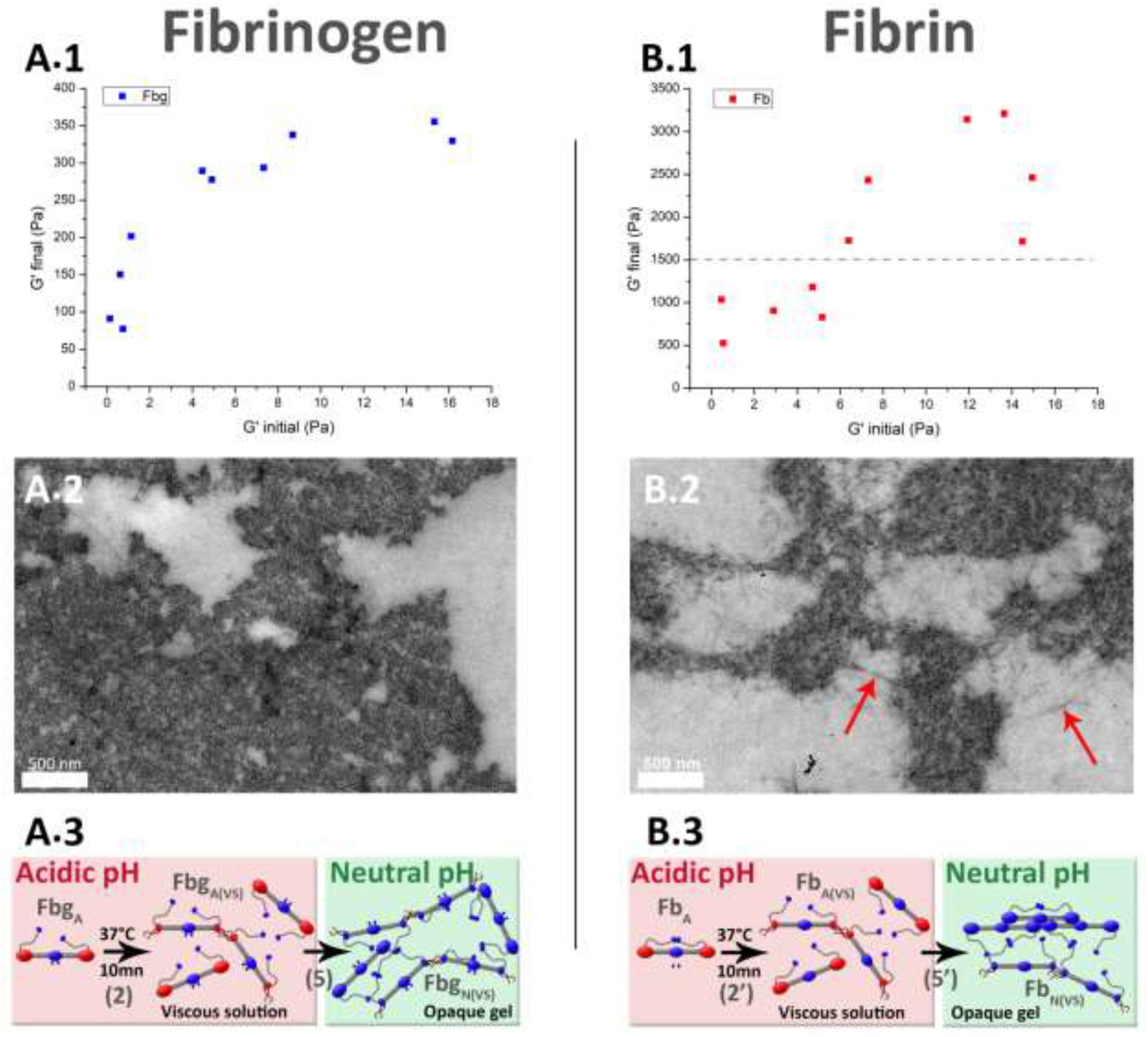
Effect of neutralization on viscous acidic solutions of fibrinogen Fbg_A(VS)_ (A) and fibrin Fb_A(VS)_ (B) and analysis of the final neutral gels Fbg_N(VS)_ and Fb_N(VS)_. (A.1 & B.1) Resulting storage moduli (G’ final) of respectively Fbg_N(VS)_ and Fb_N(VS)_, as a function of the initial storage moduli of respectively Fbg_A(VS)_ and Fb_A(VS)_ fixed by different incubation times at 37°C. (A.2 & B.2) Resulting structures, respectively Fbg_N(VS)_ and Fb_N(VS)_, as observed by TEM. (A.3 & B.3) Schematic sum up of the neutralization **(5 & 5’)** of respectively Fbg_A(VS)_ and Fb_A(VS)_, where only parts of the proteins are denatured by short incubation times **(2 & 2’)** and form aggregates. For fibrinogen Fbg_A(VS)_, upon neutralization **(5)** the strengthening of the network is hypothesized to rely on αCs interactions promoted by the proximity of proteins. For fibrin solutions Fb_A(VS)_, upon neutralization **(5’)** the non-denatured fibrin proteins selfassemble following the classical scheme, bridging aggregates formed during incubation at acidic pH **(2’)**.

#### Fibrin

The same rheological study was carried out by neutralizing acidic fibrin solutions Fb_A(VS)_ at different initial viscosities. As seen on Fig. 4.B.1, in spite of measures’ dispersity, for non incubated solutions Fb_A_ and shortly-incubated solutions Fb_A(VS)_ with low initial storage moduli, the final G’ of Fb_N_ and Fb_N(VS)_ ranged between 500 and 1250 Pa. For longer incubated Fb_A(VS)_ solutions, with higher initial G’, the final value exceeded 1500 Pa, and even 3000 Pa for certain gels. All the fibrin gels Fb_N_ and Fb_N(VS)_ reached higher storage moduli than pure fibrinogen gels Fbg_N(VS)_, independently of the initial solution viscosity. TEM observation of Fb_N(VS)_ gels exhibited a composite structure, with thin fibers connecting aggregate-like islands similar to the ones present in pure fibrinogen gels Fbg_A(G)_ and Fbg_N(VS)_ (Fig. 4.B.2). Upon neutralization, the non-denatured fibrin proteins self-assemble following the “classical” scheme, bridging the aggregates islands through thin fibers (Fig. 4.B.3). The formation of aggregates by incubation prior to neutralization has a tremendous effect on final gel properties. Indeed, for non-or shortly-incubated solutions Fb_A_ and Fb_A(VS)_ with low initial G’, the final storage modulus remains below 1.5 kPa, while viscous solutions Fb_A(VS)_ incubated for a longer time all reach higher moduli. The high dispersion of the measures may be explained by the fact that the system is more complex than for fibrinogen, with multiple mechanisms involved.

### Versatility in acidic solution viscosity and potential applications

The here presented study sheds light on the behavior of fibrinogen and fibrin at acidic pH, and the different steps involved in gelation process are depicted in Fig. 5. Beyond the interest from a biochemical standpoint the investigated phenomena represent a unique opportunity to produce stable materials made of pure fibrin (Fb_N_, Fb_N(VS)_, Fb_N(G)_ and Fb_A(G)_) and, more significantly, of pure fibrinogen (Fbg_N(VS)_, Fbg_N(G)_ and Fbg_A(G)_) without any use of chemical or physical crosslinking. Two main processes can be implemented from our work.

**Fig. 5.**
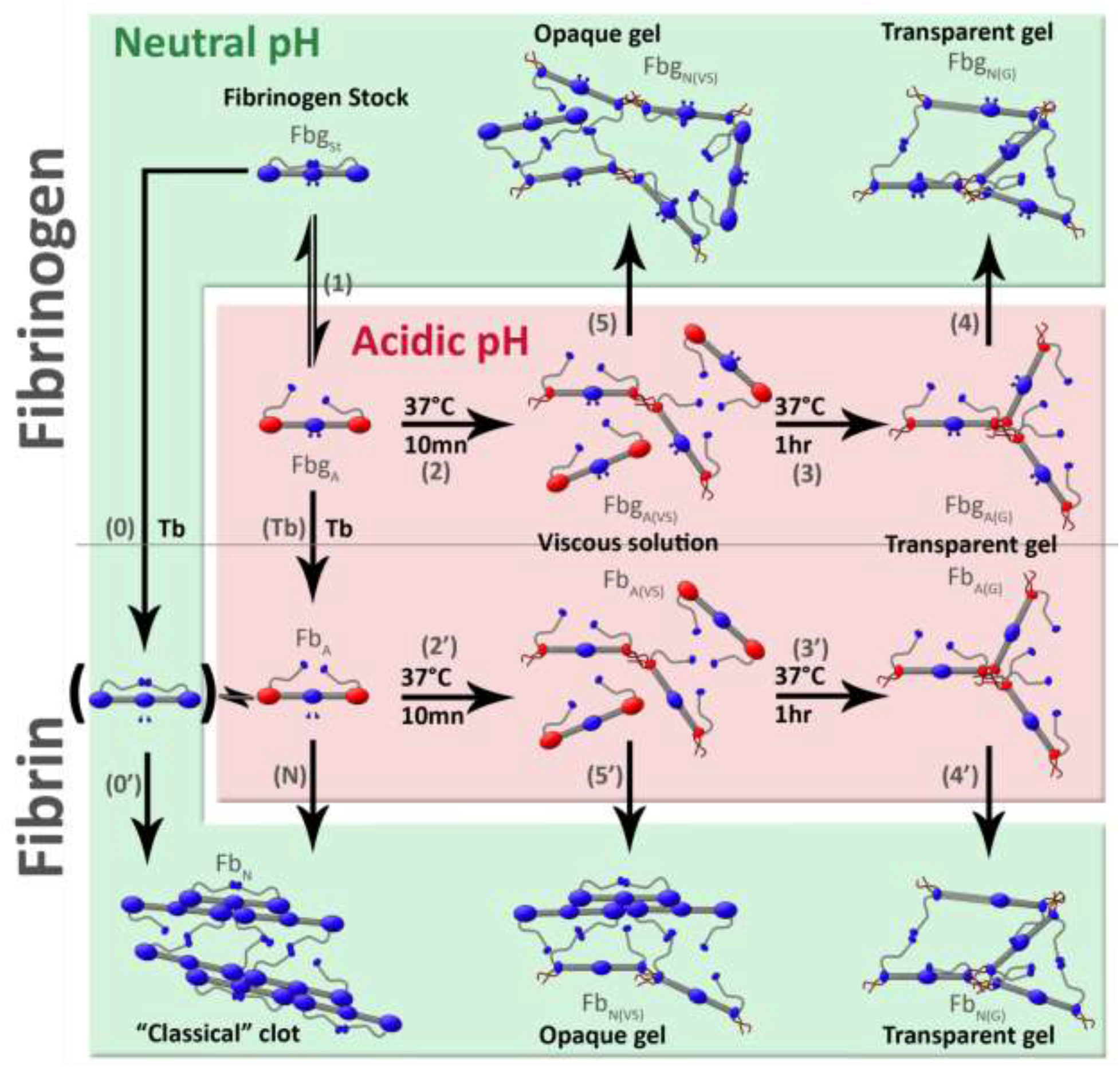
The different mechanisms of fibrin(ogen) gelation. Classically, fibrin Fb_N_ is polymerized by the action of thrombin that cleaves Fps from fibrinogen Fbg_St_ **(0)**, causing self-assembly in a half staggered manner **(0’)**. Upon acidification **(1)**, fibrinogen Fbg_St_ undergoes a reversible change in structure which involves the destabilization of the thermolabile D domains and αCs interactions in Fbg_A_. Incubation at 37°C denatures this domain, enabling the interactions of “denatured proteins” by their extremities, which causes an increase in viscosity **(2)** of the solution Fbg_A(VS)_ which eventually sets **(3)** to a gel Fbg_A(G)_. Upon neutralization, the formation **(5)** of opaque gels Fbg_N(VS)_ from viscous solutions Fbg_A(VS)_ and the strengthening **(4)** of the transparent gels Fbg_A(G)_ are hypothesized to be partly due to αCs interactions. Addition of thrombin **(Tb)** to freshly acidified fibrinogen solutions Fbg_A_ causes the cleavage of fibrinopeptides but prevents fibrin Fb_N_ assembly, which only occurs upon neutralization (N). No effect of the Fps cleavage could be observed on the Fb_A(VS)_ viscosity increase **(2’)** and gel Fb_A(G)_ formation **(3’)** of acidic solutions Fb_A_ upon incubation, compared to **(2)** and **(3)**, as well as after neutralization of Fb_A(G)_ gels **(4’)**, compared to **(4)**. However, neutralization of viscous fibrin solutions Fb_A(VS)_ **(5’)** produced composites made of acidic aggregates bridged by “classical” fibers, and resulted in stiffer gels Fb_N(VS)_, compared to those obtained (Fb_N_) from non-incubated solutions Fb_A_ after neutralization **(N)**. These phenomena are useful to produce transparent fibrinogen Fbg_N(G)_ or fibrin Fb_N(G)_ gels **(4)(4’)** and enable the production of fibrinogen Fbg_N(VS)_ **(5)** and fibrin Fb_N(VS)_ **(5’)** threads, and provide a proof of concept in printing, using incubation of solutions **(2)(2’)** to tune the solution viscosity and/or the threads final mechanical properties.

The first is the formation of stable transparent (Fbg_A(G)_ and Fbg_N(G)_) gels by incubation **(2+3)** of acidic fibrinogen solutions. Such homogeneous materials may be valuable for molding or coating processes since they are much easier to process than those obtained by mixing fibrinogen and thrombin at neutral pH **(0+0’)**. At low ionic strength, those gels do not contract and are transparent, which represents an obvious advantage for such processes. Fibrin materials (Fb_A(G)_ and Fb_N(G)_) can also be produced the same way **(2’+ 3’)**, with no obvious benefit on structural and mechanical properties as compared to those made of fibrinogen (Fig. 3.A).

The second process is the instantaneous formation of gels (Fbg_N(VS)_, Fb_N_, Fb_N(VS)_) from acidic solutions by a simple change from acidic to neutral pH **(5,5’)**, on the contrary to the delayed gelation **(0’)** initiated by thrombin **(0)** used in the literature ^23^. The pH-triggered gelation of stable acidic solutions of fibrin(ogen) enables to transpose multiple technique implemented for collagen to fibrin(ogen). For instance, this makes it possible to produce stable fibrinogen and fibrin threads (Fig. 6.A) by extrusion in neutral buffer, and provides the proof of concept for pure fibrin and fibrinogen printing as shown on Fig 6.B and SI Appendix, Video S1 where the printing of a honeycomb pattern is displayed.

**Fig. 6.**
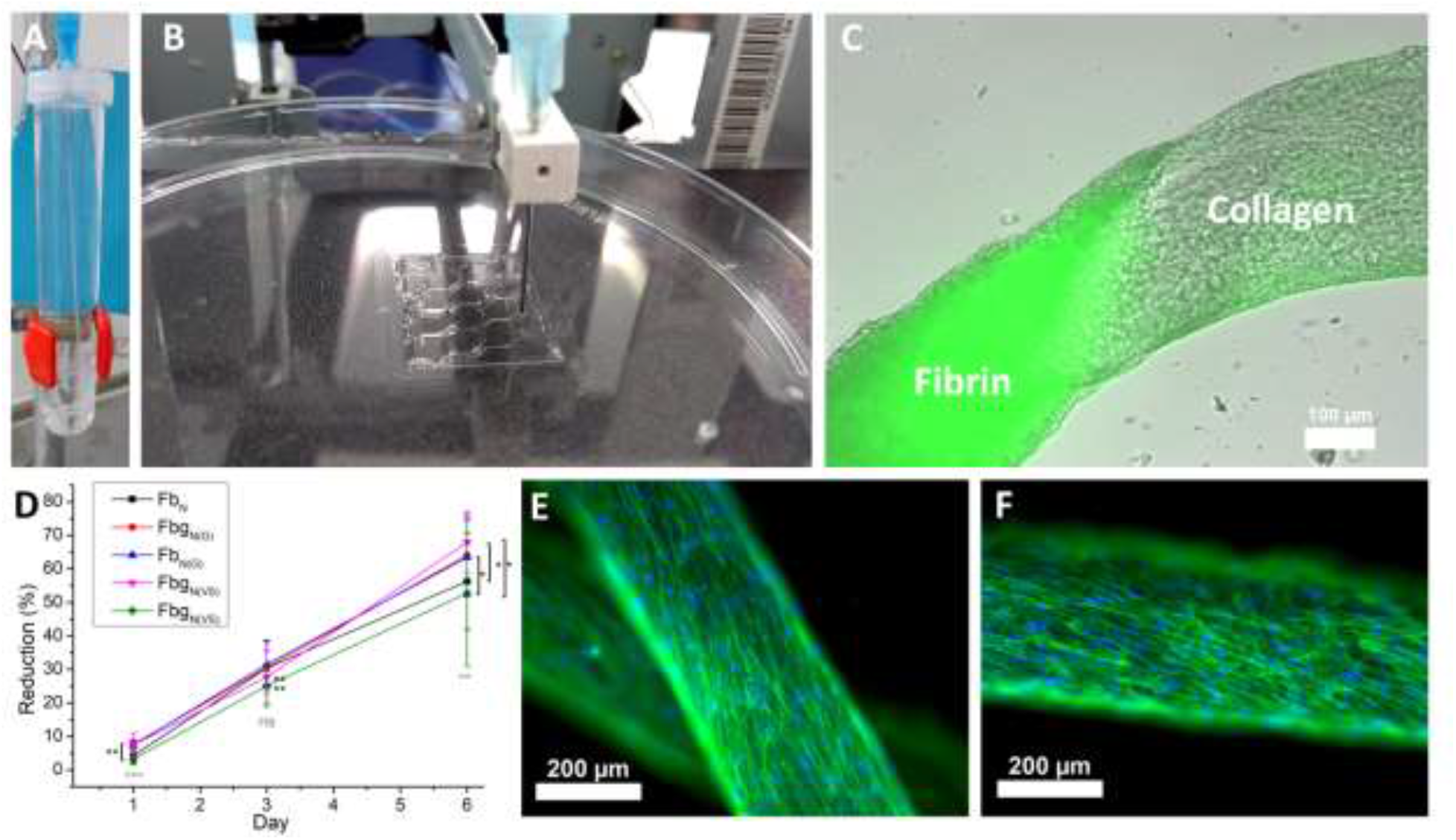
Use of acidic fibrin and fibrinogen solutions for production of new materials. (A) Image of the extrusion of a viscous fibrinogen solution in 100 mM Hepes 2.5% PEG solution pH 7.4. (B) Printing of a pure viscous fibrin solution on a Petri dish. (C) Pure fibrin to collagen gradient in a continuous thread produced by alternate extrusion of viscous acidic fibrin and collagen solutions (60 mg/mL, 3mM HCl, 0.8mM citrate pH 3.6) in the same conditions (100mM Hepes & 2.5% PEG) observed with a fluorescent microscope. Green: FITC-labelled fibrin. (D) Reduction rate of L929 murine fibroblasts seeded in wells coated with “classical” fibrin Fb_N_, acidic fibrinogen and fibrin gelled by incubation Fbg_N(G)_ and Fb_N(G)_, and viscous acidic fibrinogen and fibrin gelled by neutralization, Fbg_N(VS)_ and Fb_N(VS)_.. (E & F) Normal Human Dermal Fibroblasts 3 days after seeding on respectively a pure fibrinogen Fbg_N(VS)_ and a pure fibrin Fb_N(VS)_ thread. Blue: nuclei (DAPI); green: actin (phalloidin).

The ability to tune the viscosity **(2 or 2’)** of the solution is a key asset for processing and allows controlling the mechanical properties of the final gel (Fig. 4.A.1&B.1). Increasing solution viscosity by incubating fibrin(ogen) acidic solution has a paramount effect on thread production by extrusion, as it stabilizes the flow outside the needle. Viscosity of solutions is also a paramount parameter for printing: low viscosity fibrin solutions may be, for instance, interesting for inkjet printing, while viscous solutions are sought for extrusion printing. The acidic pH of the solutions limits the incorporation of cells directly in the inks, on the contrary to classical pathway using thrombin only. Use in bioprinting may then require to print cells and fibrin(ogen) separately. Fibrin(ogen) inks may be used as a support matrix where cells can be printed in between fibrin(ogen) threads, in the fashion of Atala’s group ^19^, or by means of co-extrusion ^44^.

Additionally, acidic conditions and extrusion are compatible with collagen solutions, enabling production of composite materials. For instance, fibrin(ogen) solution can be mixed with collagen solution (60 mg/mL, 3 mM HCl, 0.8 mM citrate, pH 3.6) at any rate from 100% fibrin(ogen) to 100% collagen and extruded in the same neutralizing buffer (100 mM Hepes, 2.5% PEG). Coextrusion can also be performed, with the possibility to tune dynamically thread composition using a double extrusion device (SI Appendix, Fig. S7) and to produce fibrin(ogen) to collagen gradients (Fig. 6.C).

Test were performed to ensure that modification of fibrin(ogen) caused by acidic incubation did not alter fibrin(ogen) biocompatibility and bioactivity. L929 murine fibroblasts, commonly used to test biocompatibility in the ISO standard, were cultured on wells coated with “classical” fibrin Fb_N_, acidic fibrinogen and fibrin gelled by incubation Fbg_N(G)_ and Fb_N(G)_, and viscous acidic fibrinogen and fibrin gelled by neutralization, Fbg_N(VS)_ and Fb_N(VS)_. As observed on Fig. 6.C, all materials induced a similar increase in metabolic activity from day 1 to day 6, due to the high proliferation of L929 on the materials till confluence (SI Appendix, Fig. S8). On all materials, cells adopted typical L929 cells morphology, demonstrating that the various processes do not prevent cell adhesion to matrix. Interestingly, at day 1 and 6, a higher metabolic rate is observed (p<0.0001 & p=0.0019 resp.) on non-self-assembled fibrin(ogen), obtained by total incubation Fbg_N(G)_ and Fb_N(G)_ or by neutralization of viscous fibrinogen solution Fbg_N(VS)_, compared to materials containing self-assembled fibrin, “classical” clot Fb_N_ and neutralized viscous fibrin solution Fb_N(VS)_.

Further test with primary human fibroblasts seeded on Fbg_N(VS)_ fibrinogen and Fb_N(VS)_ fibrin, obtained by extrusion of the corresponding solutions in a neutralizing buffer (100 mM Hepes 2.5% PEG, pH 7.4), was performed to ensure that the process and the difference of structure did not alter cell adhesion. Culture of Normal Human Dermal Fibroblasts on threads exhibited high affinity toward both types of threads, with high cellular densities (Fig. 6.E&F). After three days, both types of threads were covered with a monolayer of cells, without colonization inside the threads, as confirmed using confocal microscopy (SI Appendix, Fig. S9).

Ability to produce fibrinogen materials represents an obvious interest for biomedical applications. Indeed, it spares the need of thrombin, reducing cost and risks as thrombin used clinically is not autologous whereas fibrinogen can be directly extracted from blood patient ^45^. On a more academic point of view, ability to produce fibrinogen materials enables the investigation of their effect on cell behavior such as, for instance, the influence on cell phenotype of remaining fibrinopeptides in thrombin-free materials ^46,47^ or the presence of self-assembled fibrin.

In summary, our present results widen the possibilities in terms of fibrin processing, and enable production and shape control of fibrinogen materials. Furthermore, the processes are implemented in similar conditions as those used for collagen, which should enable the development of hybrid scaffolds based on these two proteins. The possibility to tune the biochemical and structural properties of these materials should be an asset in order to induce specific and spatially-controlled interactions with the cells and obtain appropriate cellular response to mimic the complexity of the tissues.

## Materials & Methods

### Fibrinogen and thrombin solutions

Fibrinogen stock solutions were prepared by dissolving 1 g of human fibrinogen lyophilized from 20 mM citrate pH 7.4 solutions (Merck 341576, >90% clottable proteins) into distilled water as recommended by the manufacturer (final volume of app. 22 mL). The final pH of the stock solutions was 6.6 to 6.8. The concentration (40 ± 2 mg/mL) was measured with the extinction coefficient at 280 nm (1.51 mL/mg) 27. The solutions were aliquoted and kept at −80°C. Lyophilized thrombin from bovine plasma (Sigma) was dissolved in sterile PBS 1X to reach a concentration of 200 U/mL. The solutions were aliquoted and kept at −20°C.

FITC-labeled fibrinogen was prepared by dissolution of 20 mg of FITC CeliteR in 3 mL of 0.13 M carbonate buffer pH 9.2 subsequently mixed with fibrinogen stock solution at 40 mg/mL at a 1:3 ratio and dialysed 48 hours against 2 L 20 mM citrate pH 7.4, aliquoted and stored at −80°C. For extrusion, 10% in volume of FITC-labeled fibrinogen [Fbg-FITC] was added to fibrinogen stock solution, following the same protocol as usual afterwards.

### Rheology

Rheology was performed on an MCR 302 rheometer (Anton Paar), equipped with a temperature control of both upper and lower geometries (Peltier hood PTD200). A sanded plate-plate geometry (PP25/S) was used for gel formation study and subsequent gel characterization.

For acidic gel formation study, significant amount of water was poured around the geometry, and the hood lowered to prevent from drying. Measurements were performed at constant amplitude and frequency (γ=0.1 %, ω=1 rad/s). The temperature was kept at 25°C for 5 mn and then switched to 37°C (zero-gap calibration was performed at 37°C prior to measure). Once gel formed, a frequency sweep (ω = 0.1 to 100 rad/s; γ =1 %) was performed, followed by an amplitude sweep (γ= 0.05 to 1000%; ω = 1rad/s).

Gel neutralization experiments on acidic gels were carried out following the protocols above, except that a solution of 100 mM Hepes pH 7.4 was carefully added around the geometry after 3 hours of measure, once acidic gel formation completed.

For the study of neutralization of viscous solutions a solution of 100mM Hepes, pH 7.4, was carefully added around the geometry, after 5 mn of measure. Temperature was kept constant at 25°C during the whole experiment. Frequency and amplitude sweep were performed as described above.

### Transmission electron microscopy

For inclusion in araldite, hydrated samples were crosslinked with paraformaldehyde, glutaraldehyde and osmium tetroxide 4 wt. %. They were subsequently dehydrated using baths with increasing concentrations in ethanol, progressively transferred to propylene oxide and incorporated in araldite resin prior to sectioning.

Observation of solutions was performed by dilution of solutions by a factor 10000 with the corresponding buffer and deposition of a few drops on a charged carbon coated TEM grid. The solution was sucked up after a few seconds with a filter paper on the side of the grid. Gels were chopped on a glass slide and a carbon coated TEM grid was put in contact with the mixture and dried by sucking up the supernatant with a filter paper. A drop of uranyl acetate 0.5% was then added and removed the same way.

The observations were made on a transmission electron microscope FEI Tecnai Spirit G2 operating at 120 kV. Images were recorded on a Gatan CCD camera.

### Differential Scanning Calorimetry

Differential Scanning Calorimetry was performed on a DSC Q20 from TA. Small volume (20 μL) sealed pans were used. As a reference, a similar pan with comparable weight of relevant buffer was used. Solution was kept at 20°C for 5 min for equilibration, followed by a ramp at 10°C/min till 100°C.

### Circular Dichroism

Circular Dichroism was performed on a J-810 CD from Jasco, with a temperature control Peltier module (PTC-4235). As the intensity was too high for stock solutions, even with a 0.01 mm cuvette, stock solution was diluted by a factor three with 20 mM citrate solution, pH 7.4 to reach a concentration of approx. 13 mg/mL that was further used for all the experiments. Spectra were smoothed with a Savitsky-Golay filter with a polynomial order of 2 and a smoothing window of 10 points.

### Dynamic Light Scaterring

Dynamic light scattering was performed on a Malvern Zetameter with solutions 1 mg/mL fibrinogen solutions with or without addition of D or E fragments at a ratio Fbg:fragment = 1:2 in mol. Solutions at pH 7.4 were in 20 mM citrate while acidic solutions were dialysed against 50 mM Glycine buffer and pH adjusted to 3.6.

### SDS-PAGE

Acidic fibrinogen gels were made by incubating acidic solutions of fibrinogen at 37°C overnight. They were subsequently dipped into either a 20 mM citrate solution pH 7.4 or a 8 M Urea solution. As for the controls, the same protocol was carried out using a neutral stock solution of fibrinogen and a “classical” clot made by mixing thrombin with the stock solution at neutral pH. After 4 days, the solutions in which the three different forms of fibrin(ogen) had been agitated were sampled and run on 10% SDS-PAGE gels.

### Mass spectrometry

Matrix-Assisted Laser Desorption/Ionization (MALDI) coupled with a Time Of Flight (TOF) mass spectrometer was used to study the release of the fibrinopeptides FpA and FpB, respectively 1536.6 and 1552.6 Da in mass ^48,49^, after acidic fibrinogen and fibrin gel formation. As the fibrinogen stock solution initially contained some fibrinopeptides A, the stock solution was first dialysed 24 hours against 20 mM citrate pH 7.4 with a 12-14 kDa cut-off membrane. Each gel was rinsed with mechanical agitation using 50 % acetonitrile and 50 % of the relevant buffer for each gel (20 mM citrate pH 7.4 for neutral fibrin gel, 20 mM Citrate 60 mM HCl pH 3.6 for acidic fibrin and fibrinogen gels). The rinsing solutions were measured by MALDI-TOF. As a control, a clot formed by mixing thrombin to fibrinogen stock solution at neutral pH was rinsed and both FpA and FpB could be observed at 1536.6 and 1552.6 Da respectively. The attribution of these peaks to FpA and FpB was further confirmed by MS/MS.

### Cell culture

Normal Human Dermal Fibroblasts (PromoCell™) at passage 17 were seeded at 420,000 cells/mL on fibrin and fibrinogen threads in low glucose DMEM with 10% fetal bovine serum (GIBCO). After 72 hours culture, threads were rinsed with PBS 1X and fixed with PFA 4 %. Cells were permeabilized with Triton X-100. Fluorescent labeling of the nuclei (DAPI, Invitrogen) and the actin filaments (Alexa FluorR 488 phalloidin, Invitrogen) was performed. Observations of the samples were carried out under fluorescence microscope (Axio Imager D.1, Zeiss) and confocal microscope (Leica SP5 upright Confocal).

Murine fibroblasts L929 at passage 11 were seeded to a final cell density of 2×10^4^ cell/cm^2^ in the wells of 48 well-plates coated with the different fibrin(ogen) materials. Fibrinogen solutions were acidified to reach 60 mM HCl and supplemented with 10 mM CaCl_2_ final, and 20 U/mL thrombin final for fibrin solutions or PBS 1X for fibrinogen solutions. Classical Fb_N_ coating was performed using same stock solution supplemented with same concentrations of CaCl_2_ and thrombin. Viscous fibrin(ogen) solutions were incubated as for thread production, deposited on wells and neutralized with DMEM, to produce Fbg_N(VS)_ and Fb_N(VS)_. To produce Fbg_N(G)_ and Fb_N(G)_, solutions were incubated one hour at 37°C to form a gel and neutralized by adding DMEM. Alamar blue tests were performed on 6 replicates at day 1, 3 and 6 by adding resazurin solution to fresh whole medium to reach a volume of 300 μL at a concentration of 0.01 mg/mL, incubating for 4 hours and measuring fluorescence excitation at 560 nm and emission at 590 nm using a Varioskan Lux reader from Thermo Scientific. Fluorescence imaging was performed by similar labeling as on NHDF cells and observations were done on an EVOS inverted microscope from Thermo Fisher Scientific.

## Acknowledgements

The authors would like to thank Emmanuelle Sachon, Lucrèce Matheron-Duriez and Gilles Clodic, from the Institut de Biologie de Paris Seine, for performing mass spectrometry experiments. C. R. was supported by a French Ministère de la Recherche fellowship. Other funding source: Emergence program from Sorbonne Université.

## Author contributions

C. R., T. C. and L. T. conceived the study. C. R., G. M., B. H. and L. T. performed experiments. C. R. analyzed data and developed the figures. C. R., G. M., T. C. and L. T. wrote the manuscript. All authors commented on the manuscript.

## Materials & Correspondence

Correspondence and requests for materials should be addressed to L. T. or T. C.

---

**Fig. S1:**
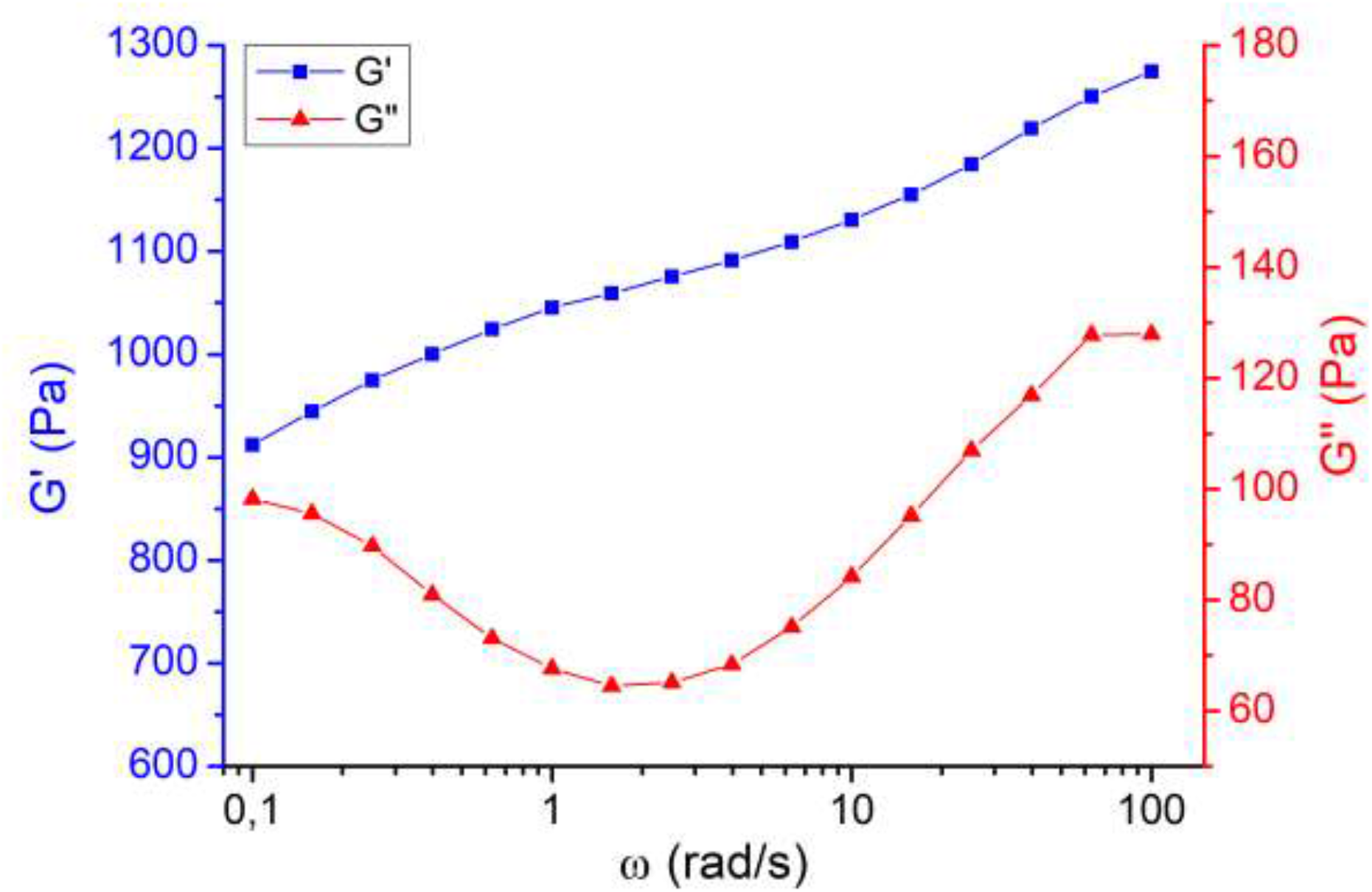
Mechanical response of an acidic fibrinogen gel Fbg_A(G)_ in frequency. Rheological study (γ=1%) of an acidic fibrinogen gel Fbg_A(G)_, polymerized prior to measure by 37°C-incubation between the geometries.

**Fig. S2:**
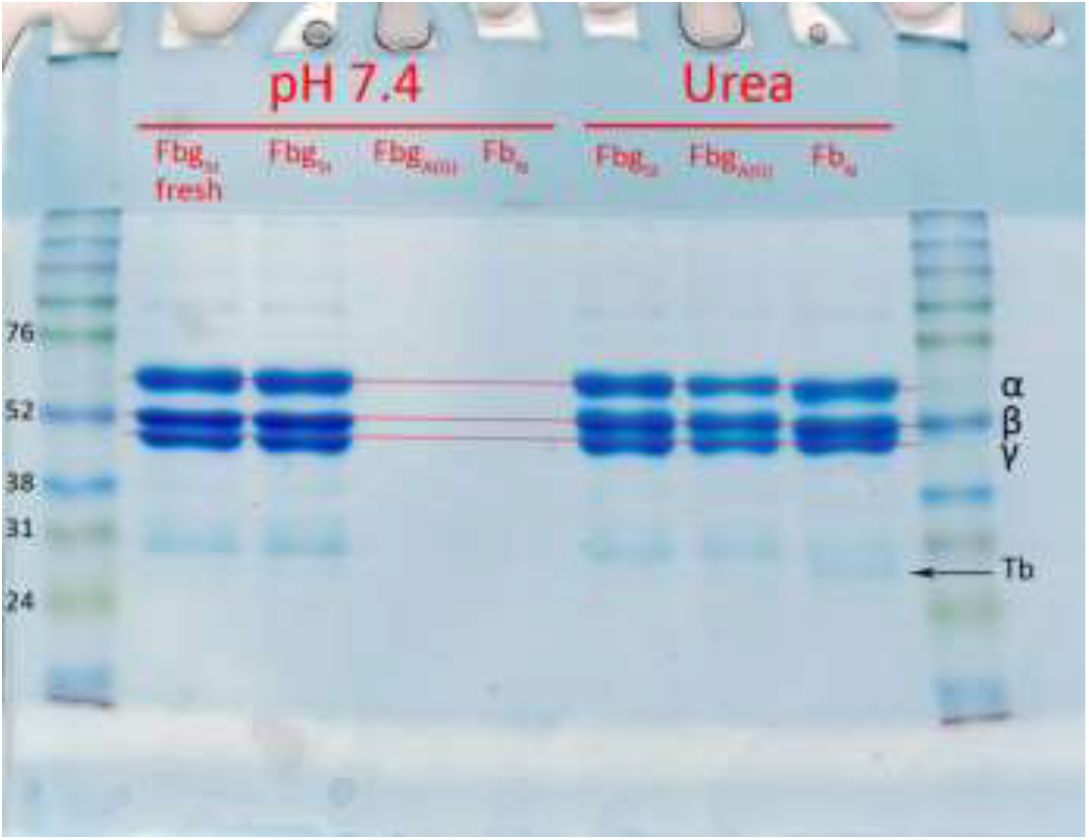
Solubility tests of acidic fibrinogen gels Fbg_A(G)_. Samples were left for 4 days in two different solutions: “pH 7.4” (20 mM citrate) and “Urea” (8 M). Supernatants were then 30ipette and run on SDS PAGE gels to investigate the presence of fibrin(ogen) proteins and possible degradation products. The samples were acid fibrinogen gels FbgA(G), neutral fibrin clots FbN and a stock solution thawed 4 days before FbgSt or the day of the SDS PAGE “FbgSt fresh”. The three main bands correspond to the three chains α, β and γ of the fibrinogen, of respectively 67, 55 and 48 kDa(1). The band around 31 kDa in FbN sample in urea corresponds to thrombin’s B chain.

**Fig. S3:**
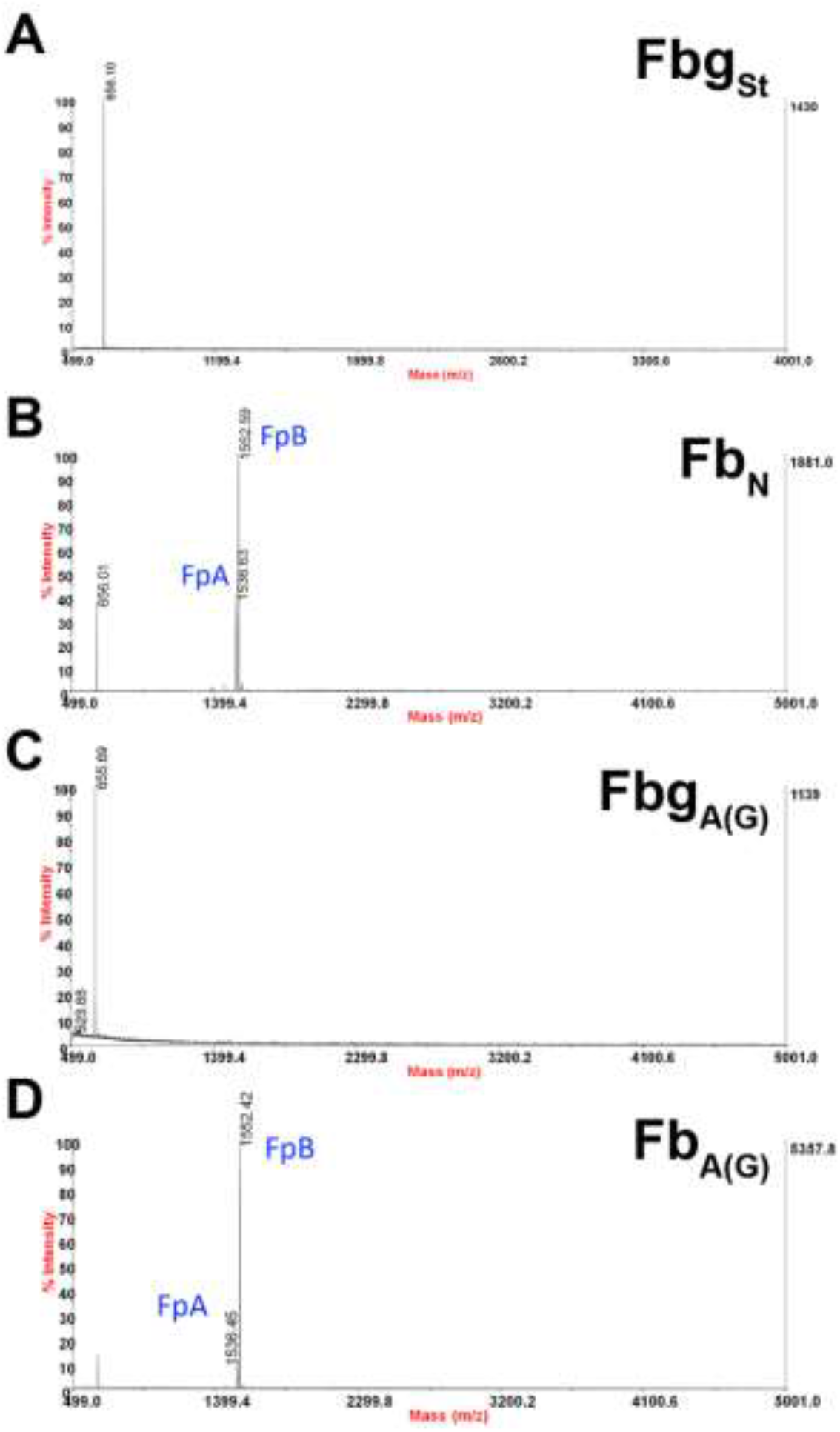
MALDI TOF of fibrinogen and fibrin in various conditions. MALDI TOF spectra of: (A) fibrinogen stock solution FbgSt (negative control), (B) supernatant after rinsing a fibrin clot FbN obtained by mixing thrombin with fibrinogen stock solution (positive control), (C) supernatant after rinsing an acidic fibrinogen gel FbgA(G) obtained after 1 hour incubation at 37°C, (D) supernatant after rinsing an acidic fibrin gel FbA(G) obtained after 1 hour incubation at 37°C.

**Fig. S4:**
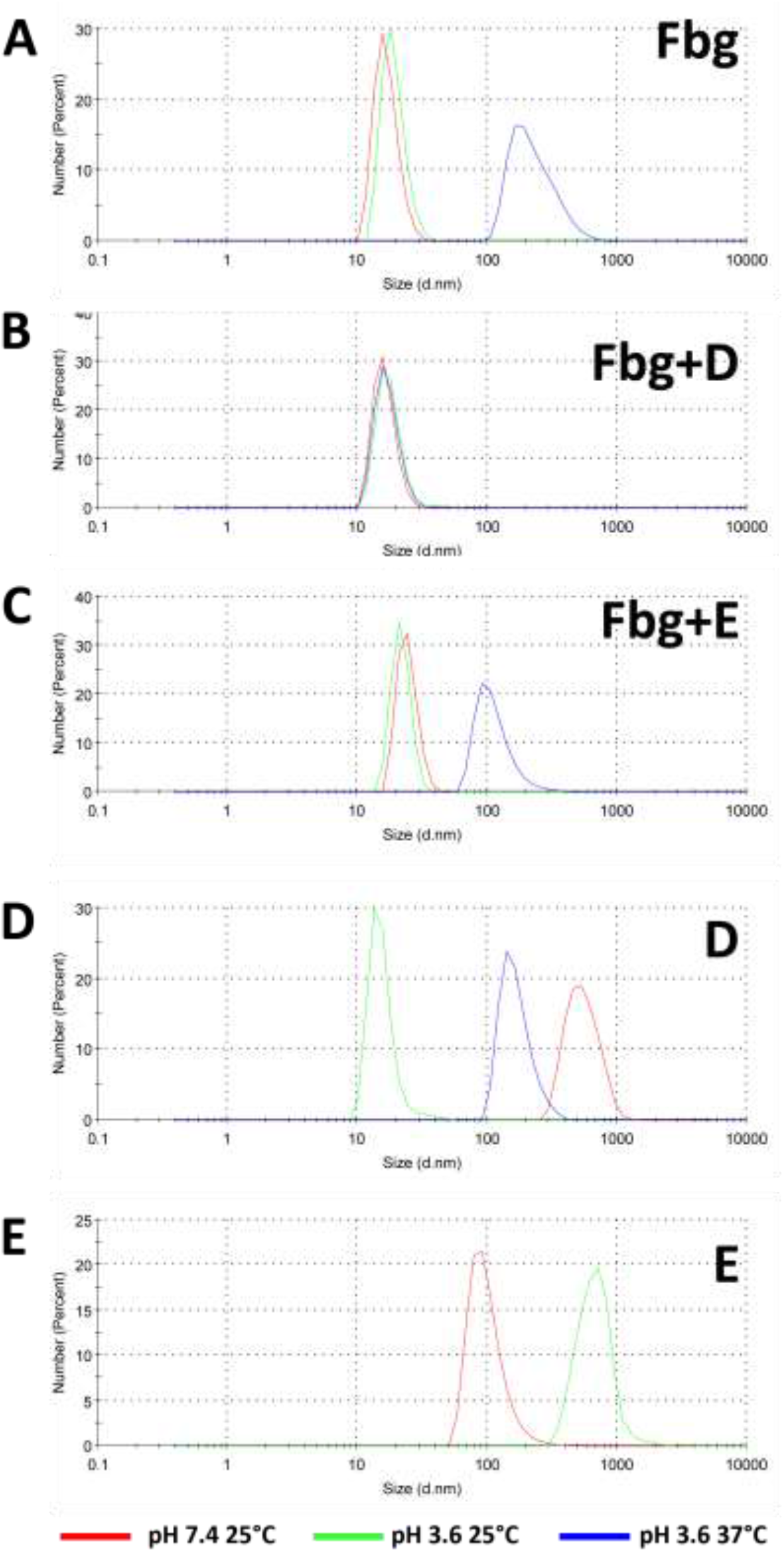
Acidic aggregation of fibrinogen with or without D or E fragments monitored by Dynamic Light Scattering (DLS). DLS counts (number) of fibrinogen (A), fibrinogen + D fragments (B), fibrinogen + E fragments (C), D fragments (D) and E fragments (E) solutions at pH 7.4 25°C (red), pH 3.6 25°C (green) and 37°C (blue).

**Fig. S5:**
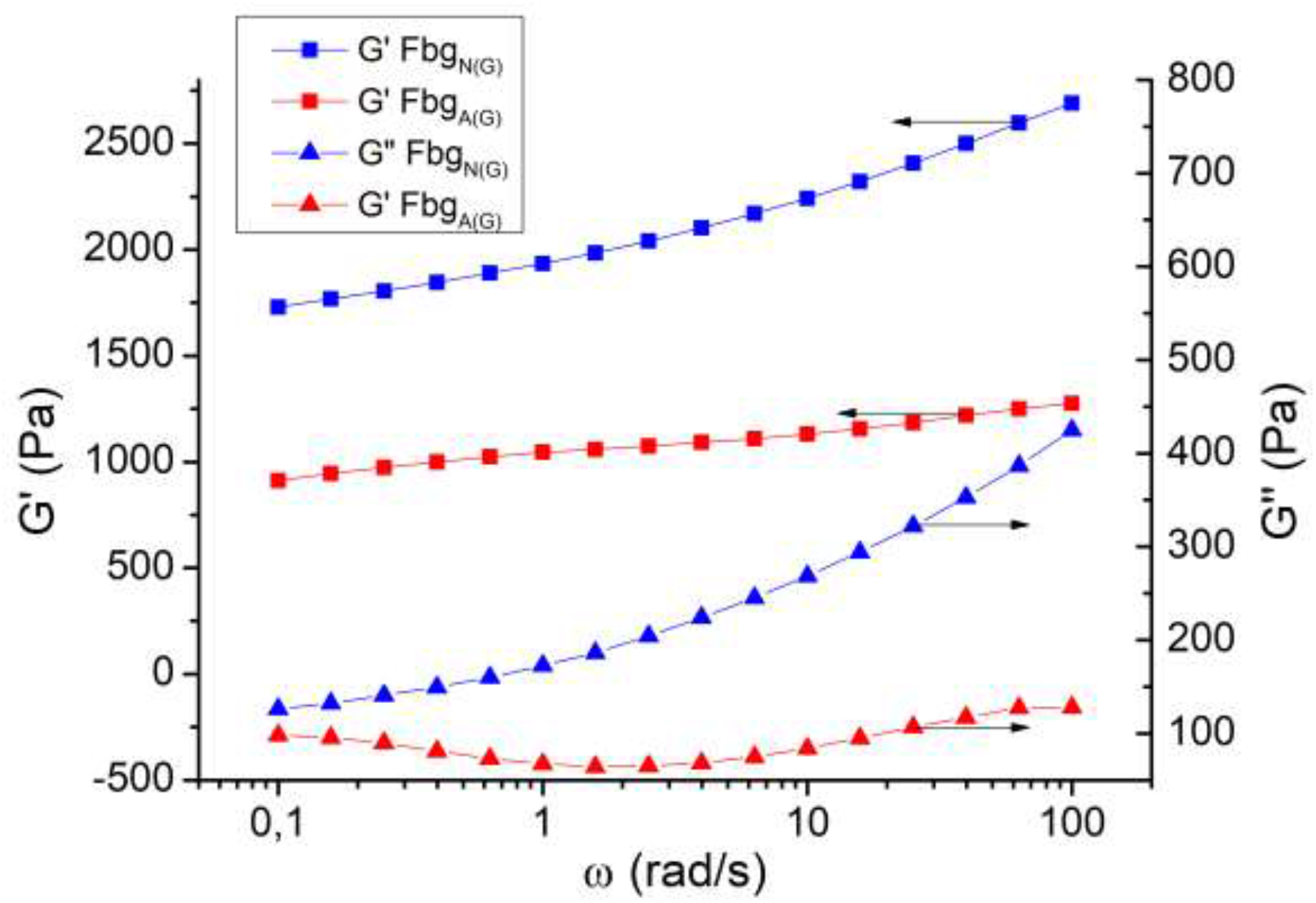
Difference in frequency between an acidic fibrinogen gel Fbg_A(G)_ and a neutralized fibrinogen gel Fbg_N(G)_. Same rheological study (γ=1%) performed on an acidic fibrinogen gel Fbg_A(G)_ (red), polymerized by 37°C-incubation between the geometries prior to measure, and a Fbg_A(G)_ gel subsequently neutralized (Fbg_N(G)_, blue).

**Fig. S6:**
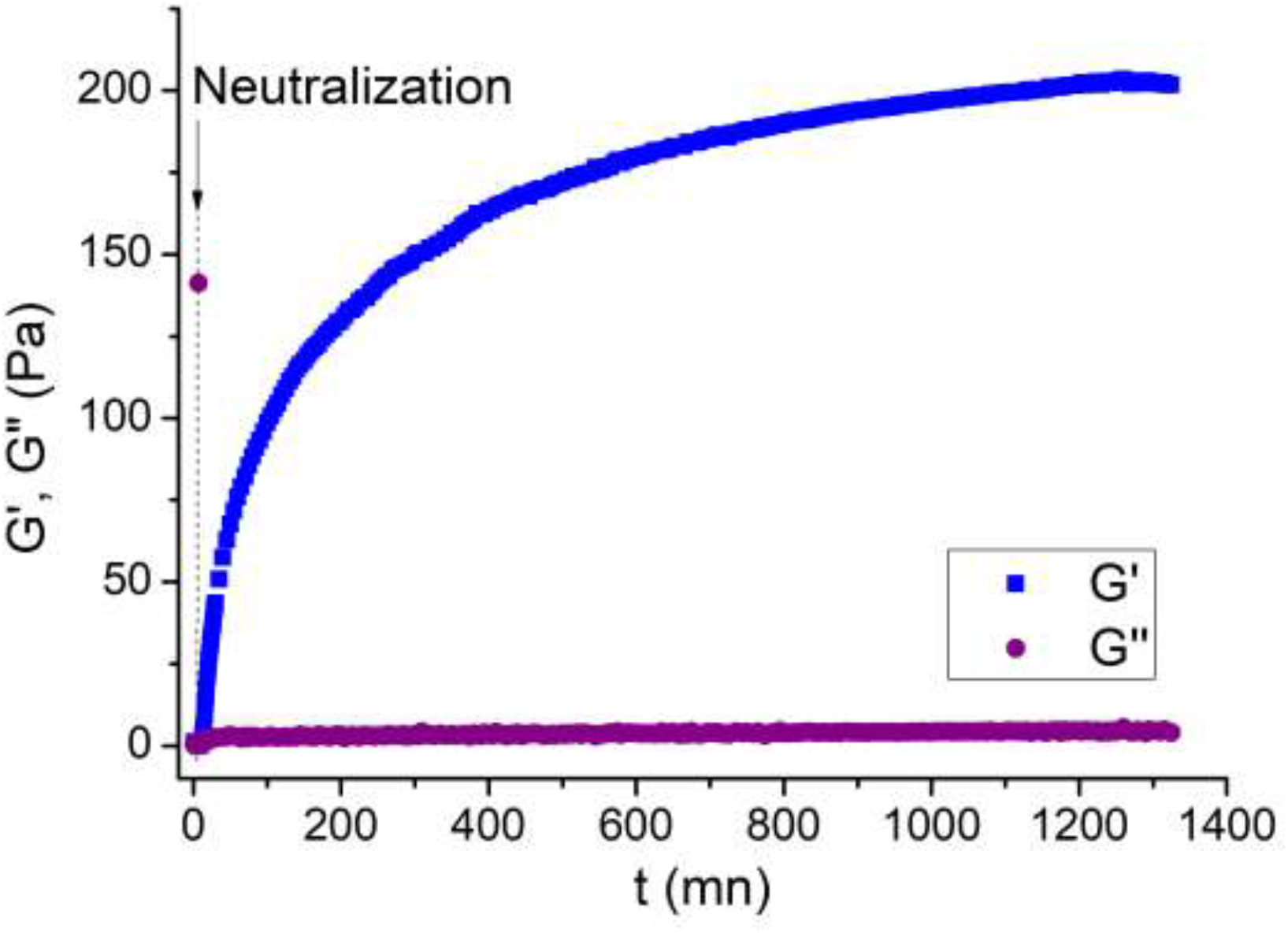
Neutralization of a viscous acidic fibrinogen solution FbgA(VS). Measurement of the rheological properties of a pre-incubated solution of acidic fibrinogen FbgA(VS) is performed. Five minutes after the start of the measure, a neutral 100 mM Hepes solution is poured around the geometry.

**Figure S7:**
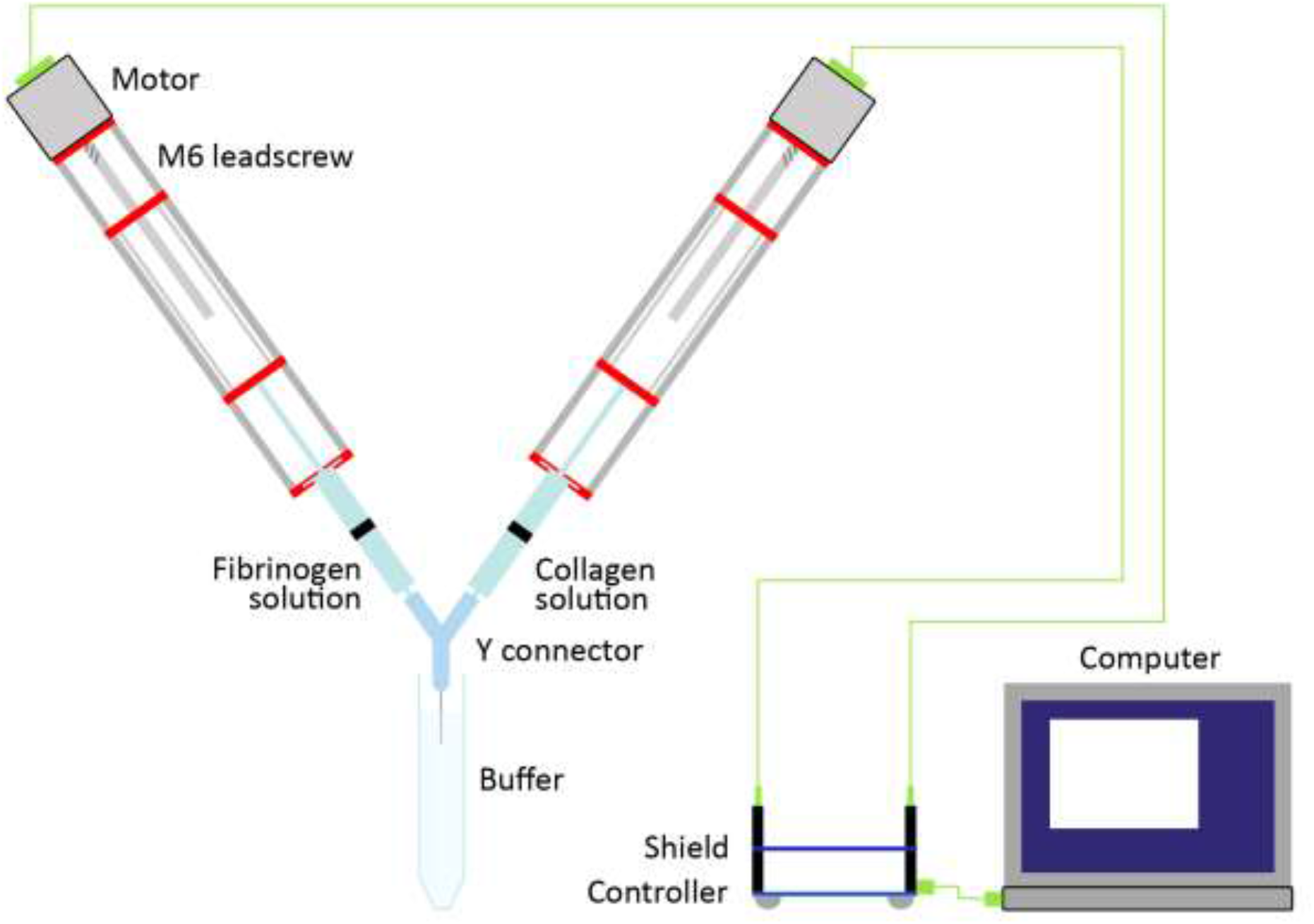
Schematic of the double extrusion device for gradient production. This lab-made device was designed to dynamically tune fibrin(ogen) and collagen content and produce collagen to fibrin(ogen) gradients thanks to controlled extrusion of both syringes thanks to a controller and to a Y connector to mix both solutions.

**Fig. S8:**
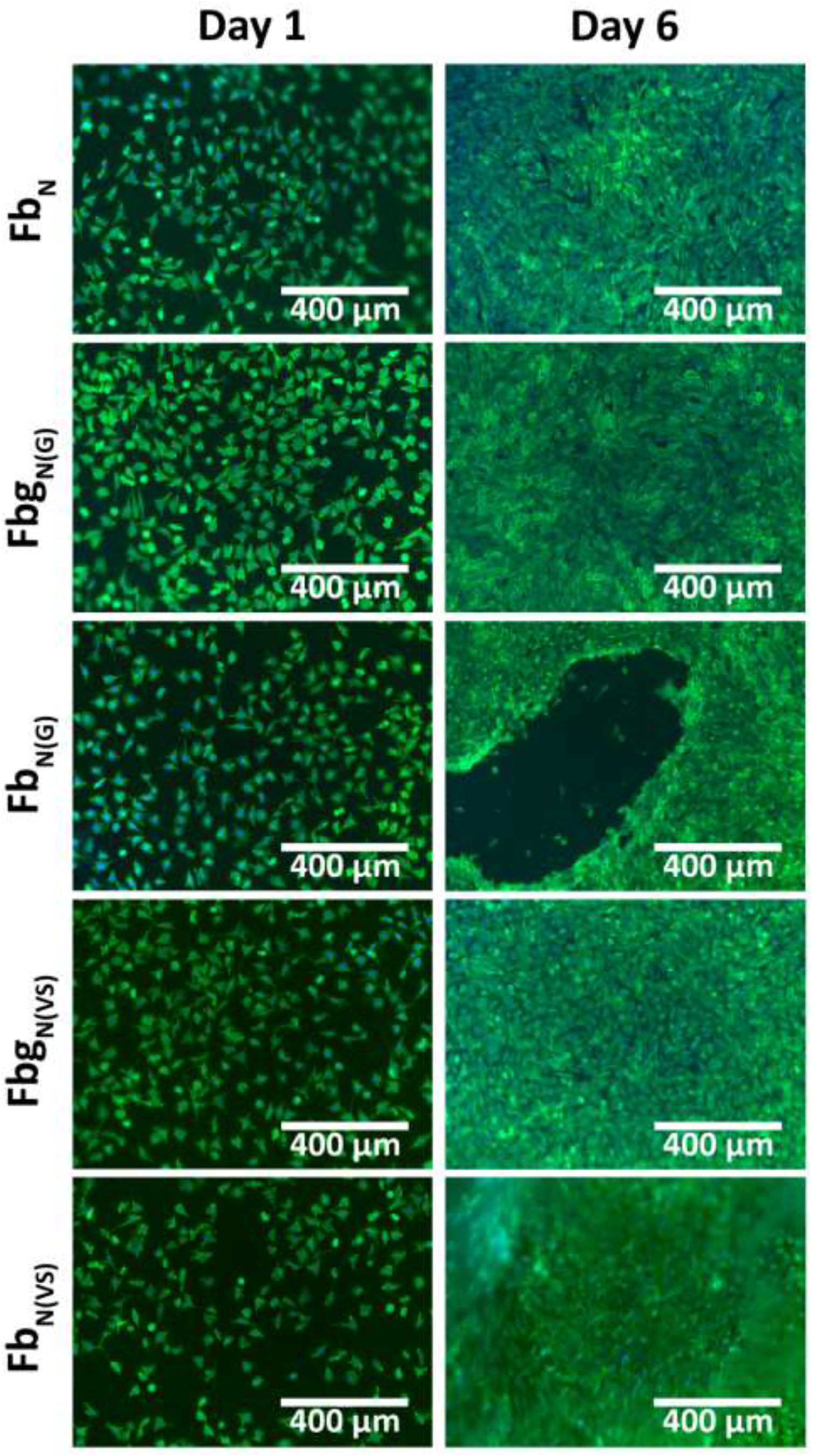
Effect of the different fibrin(ogen) materials on murine fibroblasts. Imunnofluorescence images of L929 murine fibroblasts seeded at 2×104 cells/cm^2^ in wells coated with “classical” fibrin Fb_N_, acidic fibrinogen and fibrin gelled by incubation Fbg_N(G)_ and Fb_N(G)_, and viscous acidic fibrinogen and fibrin gelled by neutralization, Fbg_N(VS)_ and Fb_N(VS)_ and observed at day 1 and 6. Blue: nuclei, Green: actin.

**Fig. S9:**
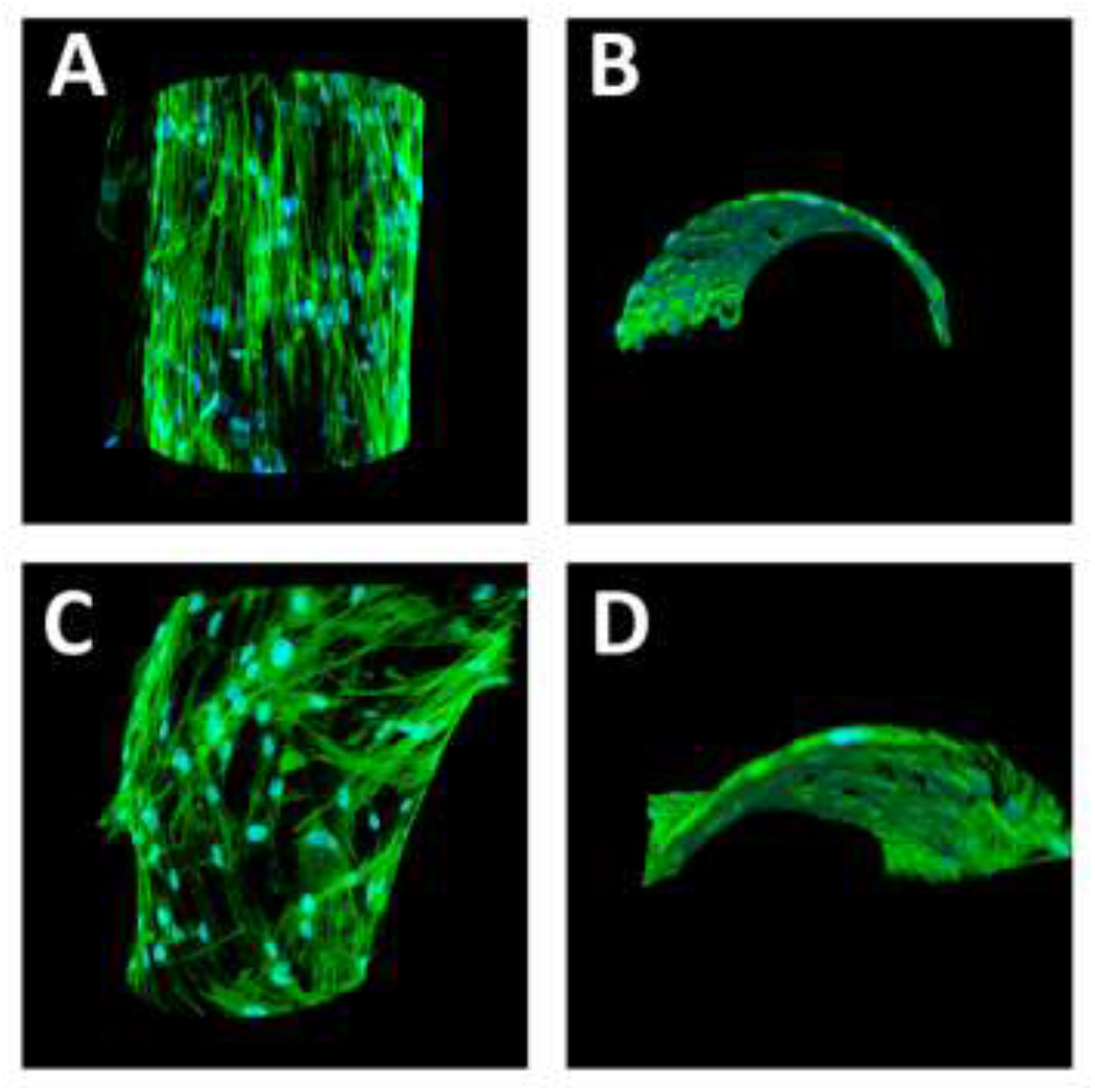
Fibrin(ogen) threads colonization by primary human fibroblasts. Confocal immunofluorescence images of NHDFs seeded on fibrin Fb_N(VS)_ (A & B) and fibrinogen Fbg_N(VS)_ (C & D) threads. Blue: nuclei, Green: actin. Dimensions: A&B 394×394×158 μm; C&D 394×394×119 μm.

**Table S1:** Solubility of acidic fibrinogen gels Fbg_A(G)_ in different solutions.

| Parameters | Solutions | Dissolution/Kinetics |
| --- | --- | --- |
| Neutral pH | 20 mM citrate pH 7.8 | No |
| Basic pH | 100 mM Tris pH 10.7 | No |
| Ionic strength | PBS 10X | No |
| Temperature | 100°C, 1h30, 20 mM citrate pH 7.8 | No |
| Weak bonds | 8 M Urea | Yes/ 10 to 15 mn |
| Disulfide bonds | 100 mM DTT pH 5/6 | Yes/ few days |
| Dilution | MQ water | No |
| Salt-in | 100 mM KSCN | No |
| Salt-in | 100 mM KI | No |
| Salt-in | 100 mM CaCl<sub>2</sub> | No |

**Video S1: Printing of a pure viscous fibrin solution in a Petri dish.**

1. Doolittle RF (2010) Fibrinogen and Fibrin. *Encyclopedia of Life Sciences*, ed John Wiley & Sons, Ltd (John Wiley & Sons, Ltd, Chichester, UK). Available at: http://doi.wiley.com/10.1002/9780470015902.a0001409.pub2 [Accessed September 16, 2016].

## References

1. Ahmed, T. A. E., Dare, E. V. & Hincke, M. Fibrin: A Versatile Scaffold for Tissue Engineering Applications. Tissue Eng. Part B Rev. 14, 199–215 (2008).

2. Janmey, P. A., Winer, J. P. & Weisel, J. W. Fibrin gels and their clinical and bioengineering applications. J. R. Soc. Interface 6, 1–10 (2009).

3. Flick, M. J. et al. Leukocyte engagement of fibrin(ogen) via the integrin receptor αMβ2/Mac-1 is critical for host inflammatory response in vivo. J. Clin. Invest. 113, 1596–1606 (2004).

4. Lishko, V. K. et al. Multiple Binding Sites in Fibrinogen for Integrin α M β 2 (Mac-1). J. Biol. Chem. 279, 44897–44906 (2004).

5. Catelas et al. Human Mesenchymal Stem Cell Proliferation and Osteogenic Differentiation in Fibrin Gels in Vitro. Tissue Eng. 12, 16 (2006).

6. Willerth, S. M., Arendas, K. J., Gottlieb, D. I. & Sakiyama-Elbert, S. E. Optimization of fibrin scaffolds for differentiation of murine embryonic stem cells into neural lineage cells. Biomaterials 27, 5990–6003 (2006).

7. Makogonenko, E., Tsurupa, G., Ingham, K. & Medved, L. Interaction of Fibrin(ogen) with Fibronectin: Further Characterization and Localization of the Fibronectin-Binding Site. Biochemistry 41, 7907–7913 (2002).

8. Weigel, P. H., Frost, S. J., LeBoeuf, R. D. & McGary, C. T. The specific interaction between fibrin(ogen) and hyaluronan: possible consequences in haemostasis, inflammation and wound healing. Ciba Found. Symp. 143, 248–261; discussion 261-264, 281–285 (1989).

9. Sahni, A., Odrljin, T. & Francis, C. W. Binding of Basic Fibroblast Growth Factor to Fibrinogen and Fibrin. J. Biol. Chem. 273, 7554–7559 (1998).

10. Semple, J. W., Italiano, J. E. & Freedman, J. Platelets and the immune continuum. Nat. Rev. Immunol. 11, 264–274 (2011).

11. Ham, S. W., Lew, W. K. & Weaver, F. A. Thrombin use in surgery: an evidence-based review of its clinical use. J. Blood Med. 1, 135 (2010).

12. Davie, E. W., Fujikawa, K. & Kisiel, W. The coagulation cascade: initiation, maintenance, and regulation. Biochemistry 30, 10363–10370 (1991).

13. Wolberg, A. S. Thrombin generation and fibrin clot structure. Blood Rev. 21, 131–142 (2007).

14. Medved, L., Weisel, J. W. & on behalf of fibrinogen and factor XIII subcomittee of the scientific standardization comittee of the international society on thrombosis and haemostasis. Recommendations for nomenclature on fibrinogen and fibrin. J. Thromb. Haemost. 7, 355–359 (2009).

15. Doolittle, R. F. Fibrinogen and Fibrin. in Encyclopedia of Life Sciences (ed. John Wiley & Sons, Ltd) (John Wiley & Sons, Ltd, 2010).

16. Scheraga, H. A. The thrombin–fibrinogen interaction. Biophys. Chem. 112, 117–130 (2004).

17. Tibbitt, M. W. & Anseth, K. S. Hydrogels as extracellular matrix mimics for 3D cell culture. Biotechnol. Bioeng. 103, 655–663 (2009).

18. Jockenhoevel, S. et al. Fibrin gel – advantages of a new scaffold in cardiovascular tissue engineering. Eur. J. Cardiothorac. Surg. 19, 424–430 (2001).

19. Kang, H.-W. et al. A 3D bioprinting system to produce human-scale tissue constructs with structural integrity. Nat. Biotechnol. 34, 312–319 (2016).

20. Gorodetsky, R. The use of fibrin based matrices and fibrin microbeads (FMB) for cell based tissue regeneration. Expert Opin. Biol. Ther. 8, 1831–1846 (2008).

21. Praveen, G., Sreerekha, P. R., Menon, D., Nair, S. V. & Chennazhi, K. P. Fibrin nanoconstructs: a novel processing method and their use as controlled delivery agents. Nanotechnology 23, 095102 (2012).

22. Rejinold, N. S. et al. Development of novel fibrinogen nanoparticles by two-step co-acervation method. Int. J. Biol. Macromol. 47, 37–43 (2010).

23. Cornwell, K. G. & Pins, G. D. Discrete crosslinked fibrin microthread scaffolds for tissue regeneration. J. Biomed. Mater. Res. A 82A, 104–112 (2007).

24. Picaut, L. et al. Pure dense collagen threads from extrusion to fibrillogenesis stability. Biomed. Phys. Eng. Express 4, 035008 (2018).

25. Rieu, C., Picaut, L., Mosser, G. & Trichet, L. From Tendon Injury to Collagen-based Tendon Regeneration: Overview and Recent Advances. Curr. Pharm. Des. 3483–3506 (2017).

26. Fay, M. & Hendrix, B. M. The effect of acid denaturation upon the combining power of fibrinogen. J. Biol. Chem. 93, 667–675 (1931).

27. Mihalyi, E. Physicochemical studies of bovine fibrinogen : III. Optical rotation of the native and denatured molecule. Biochim. Biophys. Acta 102, 487–499 (1965).

28. Wufsus, A. R. et al. Elastic Behavior and Platelet Retraction in Low- and High-Density Fibrin Gels. Biophys. J. 108, 173–183 (2015).

29. Piechocka, I. K. et al. Multi-scale strain-stiffening of semiflexible bundle networks. Soft Matter 12, 2145–2156 (2016).

30. Litvinov, R. I. & Weisel, J. W. Fibrin mechanical properties and their structural origins. Matrix Biol. (2016). doi:10.1016/j.matbio.2016.08.003

31. Liu, C., He, J., Ruymbeke, E. van, Keunings, R. & Bailly, C. Evaluation of different methods for the determination of the plateau modulus and the entanglement molecular weight. Polymer 47, 4461–4479 (2006).

32. Lim, B. B. C., Lee, E. H., Sotomayor, M. & Schulten, K. Molecular Basis of Fibrin Clot Elasticity. Structure 16, 449–459 (2008).

33. Zhmurov, A. et al. Mechanism of Fibrin(ogen) Forced Unfolding. Structure 19, 1615–1624 (2011).

34. Veklich, Y., Gorkun, O. V., Medved, L., Nieuwenhuizen & Weisel, J. W. Carboxyl-terminal Portions of the Alpha Chains of Fibrinogen and Fibrin. 268, 13577–13585 (1993).

35. Chen, Y., Mao, H., Zhang, X., Gong, Y. & Zhao, N. Thermal conformational changes of bovine fibrinogen by differential scanning calorimetry and circular dichroism. Int. J. Biol. Macromol. 26, 129–134 (1999).

36. Upchurch Jr, G. R., Ramdev, N., Walsh, M. T. & Loscalzo, J. Prothrombotic consequences of the oxidation of fibrinogen and their inhibition by aspirin. J. Thromb. Thrombolysis 5, 9–14 (1998).

37. Budzynski, A. Z. Difference in conformation of fibrinogen degradation products as revealed by hydrogen exchange and spectropolarimetry. Biochim. Biophys. Acta BBA - Protein Struct. 229, 663–671 (1971).

38. Medved, L. V., Privalov, P. L. & Ugarova, T. P. Isolation of thermostable structure from the fibrinogen D fragment. FEBS Lett. 146, 339–342 (1982).

39. Fan, F. & Mayo, K. H. Effect of pH on the Conformation and Backbone Dynamics of a 27-Residue Peptide in Trifluoroethanol: an NMR and CD study. J. Biol. Chem. 270, 24693–24701 (1995).

40. Privalov, P. L. & Medved, L. V. Domains in the Fibrinogen Molecule. J. Mol. Biol. 159, 665–683 (1982).

41. Medved, L. V., Gorkun, O. V. & Privalov, P. L. Structural organization of C-terminal parts of fibrinogen Aα-chains. FEBS Lett. 160, 291–295 (1983).

42. Medved’, L. V., Litvinovich, S. V. & Privalov, P. L. Domain organization of the terminal parts in the fibrinogen molecule. FEBS Lett. 202, 298–302 (1986).

43. Laki, K. The polymerization of proteins : the action of thrombin on fibrinogen. J. Biol. Chem. 222, 815–821 (1951).

44. Yeo, M., Lee, J.-S., Chun, W. & Kim, G. H. An Innovative Collagen-Based Cell-Printing Method for Obtaining Human Adipose Stem Cell-Laden Structures Consisting of Core–Sheath Structures for Tissue Engineering. Biomacromolecules 17, 1365–1375 (2016).

45. Spotnitz, W. D. Fibrin Sealant: The Only Approved Hemostat, Sealant, and Adhesive—a Laboratory and Clinical Perspective. International Scholarly Research Notices (2014). doi: 10.1155/2014/203943

46. Senior, R. M., Skogen, W. F., Griffin, G. L. & Wilner, G. D. Effects of fibrinogen derivatives upon the inflammatory response. Studies with human fibrinopeptide B. J. Clin. Invest. 77, 1014–1019 (1986).

47. Gray, A. J., Reeves, J. T., Harrison, N. K., Winlove, P. & Laurent, G. J. Growth factors for human fibroblasts in the solute remaining after clot formation. 4 (1990).

48. Jai-Nhuknan, J. & Cassady, C. J. Negative ion matrix-assisted laser desorption/ionization time-of-flight post-source decay calibration by using fibrinopeptide B. J. Am. Soc. Mass Spectrom. 9, 540–544 (1998).

49. Dashtiev, M. et al. Positive and negative analyte ion yield in matrix-assisted laser desorption/ionization. Int. J. Mass Spectrom. 268, 122–130 (2007).

